# The SP140-RESIST pathway regulates interferon mRNA stability and antiviral immunity

**DOI:** 10.1101/2024.08.28.610186

**Authors:** Kristen C. Witt, Adam Dziulko, Joohyun An, Filip Pekovic, Arthur Xiuyuan Cheng, Grace Y. Liu, Ophelia Vosshall Lee, David J. Turner, Azra Lari, Moritz M. Gaidt, Roberto Chavez, Stefan A. Fattinger, Preethy Abraham, Harmandeep Dhaliwal, Angus Y. Lee, Dmitri I. Kotov, Laurent Coscoy, Britt A. Glaunsinger, Eugene Valkov, Edward B. Chuong, Russell E. Vance

**Author notes:** Current address: Research Institute of Molecular Pathology, Vienna BioCenter, Vienna, Austria (Moritz M. Gaidt).

## Abstract

Type I interferons (IFN-Is) are essential for antiviral immunity but must be tightly regulated^1–3^. The conserved transcriptional repressor SP140 inhibits interferon beta (*Ifnb1*) expression via an unknown mechanism^4,5^. Here we report that SP140 does not directly repress *Ifnb1* transcription. Instead, SP140 negatively regulates *Ifnb1* mRNA stability by directly repressing the expression of a previously uncharacterized regulator we call RESIST (REgulated Stimulator of Interferon via Stabilization of Transcript, previously annotated as Annexin-2 Receptor). RESIST promotes *Ifnb1* mRNA stability by counteracting *Ifnb1* mRNA destabilization mediated by the Tristetraprolin (TTP) family of RNA-binding proteins and the CCR4-NOT deadenylase complex. SP140 localizes within nuclear bodies, punctate structures that play important roles in silencing DNA virus gene expression in the nucleus^4^. Consistent with this observation, we found that SP140 inhibits replication of the gammaherpesvirus MHV68. The antiviral activity of SP140 was independent of its ability to regulate *Ifnb1*. Our results establish dual antiviral and interferon regulatory functions for SP140. We propose that SP140 and RESIST participate in antiviral effector-triggered immunity^6,7^.

## Main

Type I interferons (IFN-Is) are cytokines that play central roles in antiviral immunity, autoimmunity, and cancer^1,2,8^. IFN-Is include *Ifnb1* and numerous *Ifna* and other isoforms, which signal through the IFN alpha receptor (IFNAR) to induce hundreds of Interferon Stimulated Genes (ISGs) that counter infection^8^. The pathways leading to *Ifnb1* induction have been enumerated in detail^3,9^. However, despite a longstanding appreciation that *Ifnb1* mRNAs turn over rapidly in cells^10^, relatively little is known about the pathways controlling *Ifnb1* mRNA stability. This is surprising given that mRNA turnover is a critical point of regulation for many other cytokines^11,12^. In addition, substantial evidence points to the importance of negative regulation of IFN-I, as excessive IFN-I can drive autoimmunity^1^ and susceptibility to bacterial infections^3^.

SP140 is an evolutionarily conserved but poorly characterized member of the Speckled Protein (SP) family of epigenetic readers that contain histone-recognition and/or DNA-binding domains, as well as an oligomerization domain structurally homologous to caspase-activation and recruitment domains (CARDs)^4^. The SP CARD mediates formation of nuclear bodies (NBs)^13^, large complexes that orchestrate various nuclear functions including transcriptional regulation^4^. The functions of SP140 are unclear, although previous work suggests SP140 is a transcriptional repressor essential for macrophage and possibly T-cell function^4,14-21^. In humans, loss-of-function mutations in SP140 are associated with immune disorders such as multiple sclerosis and B cell cancers^22–27^. We previously found a critical role for SP140 in the repression of IFN-I *in vivo*, as *Sp140^−/−^*mice are highly susceptible to multiple bacterial infections due to elevated IFN-I^5^. However, the mechanism by which SP140 represses IFN-I is entirely unknown.

### SP140 inhibits *Ifnb1* mRNA stability

To investigate how SP140 negatively regulates IFN-I, we first characterized *Ifnb1* transcript levels in wild-type C57BL/6J (B6) and *Sp140^−/−^*bone marrow-derived macrophages (BMMs) stimulated with agonists of distinct IFN-I-inducing pathways, including bacterial lipopolysaccharide (LPS), the dsRNA mimic poly(I:C), and the mouse STING agonist DMXAA (Fig. 1a). We noted that *Ifnb1* transcript levels were similar between B6 and *Sp140^−/−^*cells at early timepoints (2-4 hours) after stimulation, but remained elevated (∼10-fold) in *Sp140^−/−^* cells at late time points (8 hours) for all IFN-I inducing stimuli (Fig. 1a). A time course of *Ifnb1* transcript levels after DMXAA stimulation confirmed that *Sp140^−/−^* cells displayed increased *Ifnb1* mRNA at late timepoints after stimulation (8-12 hours) (Fig. 1b).

**Figure 1.**
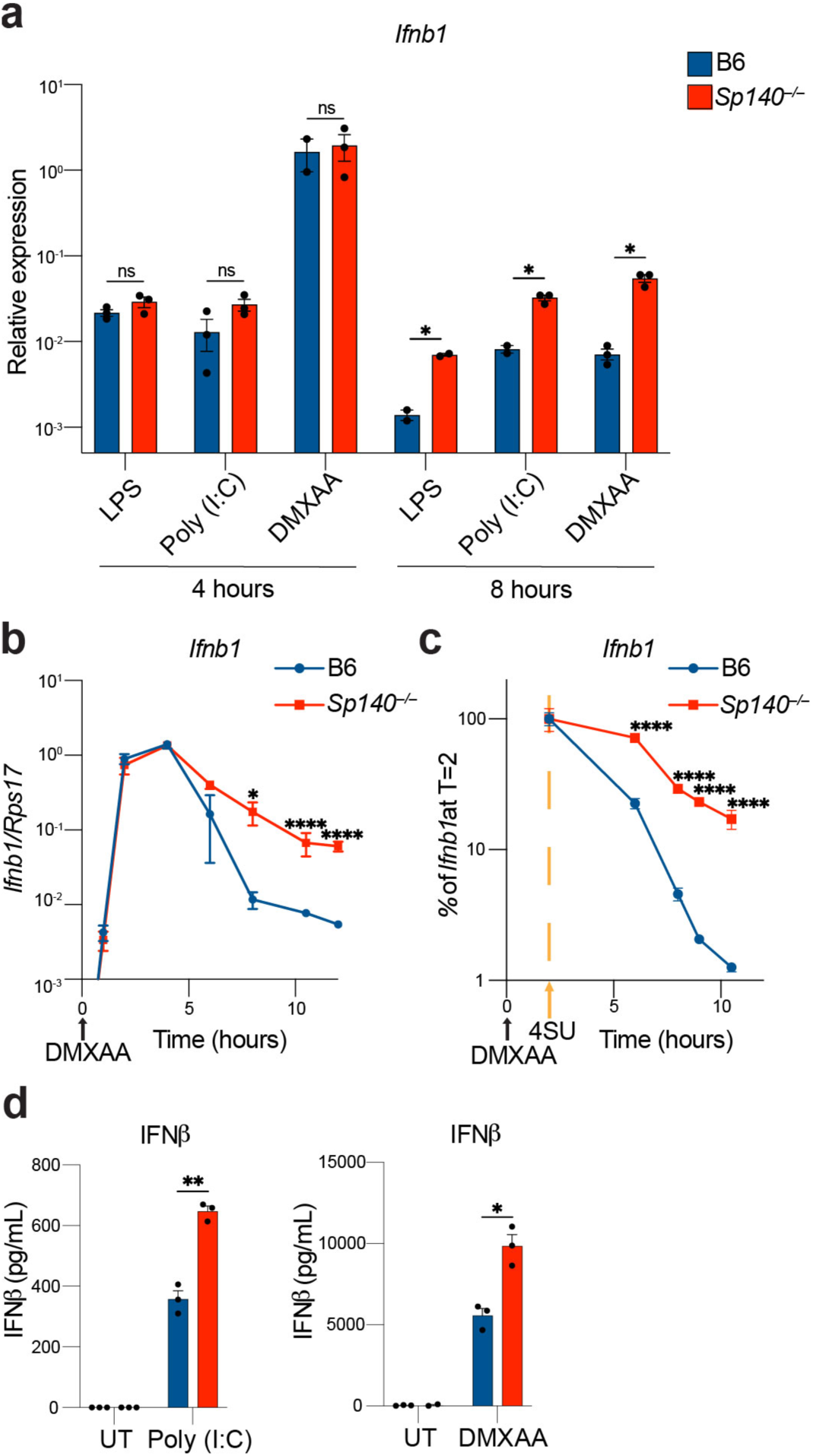
*Ifnb1* mRNA is stabilized in the absence of SP140. a. *Ifnb1* RT-qPCR in BMMs treated for 4 or 8 hours with 10 ng/mL LPS, or 100 μg/mL poly(I:C) or DMXAA. *p* values (* = *p* < 0.05, ** = *p* < 0.005, *** = *p* < 0.0005, **** = *p* < 0.00005, ns = not significant), calculated with two-tailed t-tests using Welch’s correction and FDR correction. b. *Ifnb1* RT-qPCR from BMMs treated with 100 μg/mL DMXAA at indicated timepoints. *p* values (* = *p* < 0.05, ** = *p* < 0.005, *** = *p* < 0.0005, **** = *p* < 0.00005, ns = not significant) calculated with a two-way ANOVA and Šidák’s multiple comparison correction. c. Roadblock RT-qPCR of BMMs treated with 4SU 2 hours after treatment with 100 μg/mL DMXAA. *p* values (* = *p* < 0.05, ** = *p* < 0.005, *** = *p* < 0.0005, **** = *p* < 0.00005, ns = not significant) calculated with a two-way ANOVA and Šidák’s multiple comparison correction. d. ELISA for IFNβ protein levels in supernatants of BMMs treated for 24 hours with 100 μg/mL poly(I:C) or DMXAA. *p* values (* = *p* < 0.05, ** = *p* < 0.005, *** = *p* < 0.0005, **** = *p* < 0.00005, ns = not significant), calculated with two-tailed t-tests using Welch’s correction.

We hypothesized that the elevated levels of *Ifnb1* mRNA at late time points in *Sp140^−/−^* BMMs are due to the increased stability of *Ifnb1* transcripts. To assess *Ifnb1* mRNA stability, we used Roadblock RT-qPCR^28^, a method in which the nucleotide analog 4-thiouridine (4SU) is added to cells and incorporated into newly transcribed mRNAs. Isolated RNA is then treated with N-ethylmaleimide (NEM), which reacts with 4SU to introduce a sterically bulky group that blocks reverse transcription. By adding 4SU at the peak of *Ifnb1* induction (t=2h after DMXAA stimulation), we could follow the subsequent decay of existing *Ifnb1* mRNA without detecting newly transcribed mRNAs. Using this approach, we found that the decay of *Ifnb1* transcript was markedly delayed in *Sp140^−/−^* BMMs (Fig. 1c). The increased stability of *Ifnb1* transcript in *Sp140^−/−^* BMMs resulted in an increase in IFNβ protein levels in culture supernatants (Fig. 1d). These results demonstrate an unexpected role for SP140 in the regulation of *Ifnb1* transcript stability.

### SP140 represses RESIST

SP140 lacks predicted RNA-binding domains and is believed to act in the nucleus to repress transcription^4^. We therefore hypothesized that SP140 indirectly regulates *Ifnb1* mRNA stability by repressing the transcription of an unknown factor. To identify this factor, we generated RNA-seq data from DMXAA-treated B6 and *Sp140^−/−^* BMMs, and from DMXAA-treated *Ifnar^−/−^* and *Sp140^−/−^Ifnar^−/−^*BMMs. The latter dataset eliminates the potentially confounding effects of elevated IFN-I signaling in *Sp140^−/−^* BMMs. Surprisingly, few genes were differentially expressed between DMXAA-treated *Ifnar^−/−^*and *Sp140^−/−^Ifnar^−/−^*BMMs (Fig. 2a). Other than *Ifnb1* itself, no known IFN-I regulators were differentially expressed, and gene ontology analysis of differentially expressed genes (DEGs) did not identify significantly enriched biological processes. Interestingly, only two DEGs correlated with *Ifnb1* upregulation across both RNA-seq datasets (Fig. 2b): *Sp140,* which was downregulated in SP140-deficient cells, as expected, and *Gm21188*, a poorly annotated gene that was upregulated in SP140-deficient cells. *Gm36079,* a tandem paralog of *Gm21188,* was upregulated in both datasets, but only significantly so in IFNAR-sufficient *Sp140^−/−^*BMMs treated with DMXAA. The open reading frames of *Gm21188* and *Gm36079* differ by a single silent nucleotide substitution and thus encode an identical 20.9 kDa protein. Based on our results below, we refer to this protein as RESIST (<u>RE</u>gulated <u>S</u>timulator of Interferon via <u>S</u>tabilization of <u>T</u>ranscript) and the *Gm21188* and *Gm36079* genes as *Resist1* and *Resist2*, respectively

**Figure 2.**
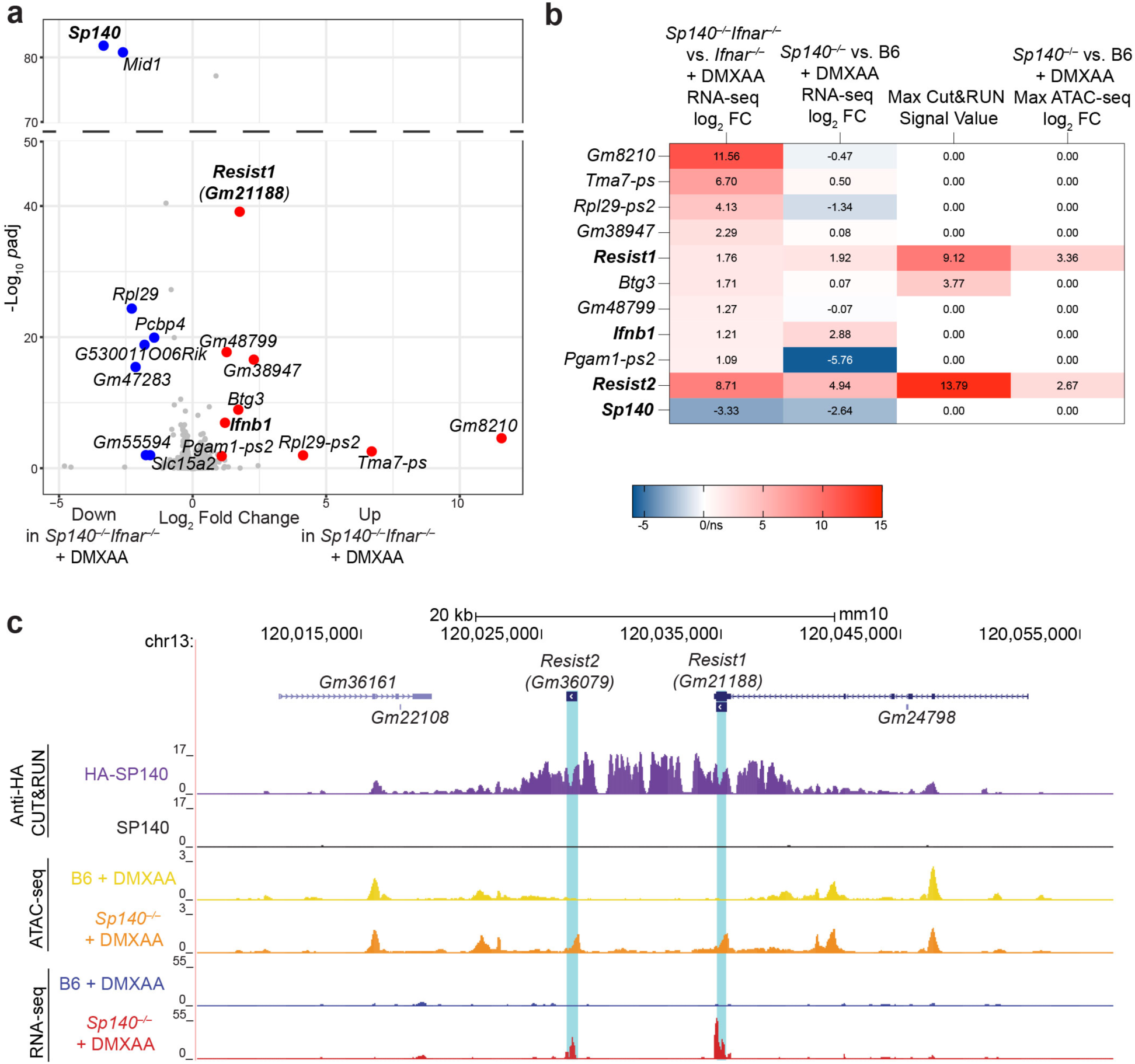
*Resist1 (Gm21188)* and *Resist2 (Gm36079)* are repressed by SP140 and correlate with increased *Ifnb1* transcript in *Sp140^−/−^* cells. a. Volcano plot of differentially expressed genes (DEGs) from RNA-seq of DMXAA-treated *Sp140^−/−^Ifnar^−/−^* vs. *Ifnar^−/−^* BMMs. Red genes are upregulated in *Sp140^−/−^ Ifnar^−/−^* BMMs with log_2_ fold change > 1 and adjusted *p-*value (padj) < 0.05. Blue genes are downregulated in *Sp140^−/−^Ifnar^−/−^* BMMs with log_2_ fold change > –1 and adjusted *p-*value < 0.05. The adjusted *p*-value for *Sp140* is < 2.225074 e-308 and is graphed as –10*(adjusted *p-*value) of *Mid1* for visualization. *Resist2* (*Gm36079)* is not depicted on volcano plot as it is removed by the DeSeq2 independent filtering function for genes with low read counts. b. Table of maximum HA-SP140 CUT&RUN MACS2 signal value, maximum log_2_ fold change in chromatin accessibility from ATAC-seq of DMXAA-treated B6 and *Sp140^−/−^* BMMs, and log_2_ fold change from RNA-seq of DMXAA-treated B6, *Sp140^−/−^, Ifnar^−/−^*and *Sp140^−/−^Ifnar^−/−^* BMMs, for significantly upregulated DEGs from a, as well as *Sp140* and *Resist2* (*Gm36079)* (ns = not significant). Cells are colored according to the column value. c. Alignment of reads at *Resist1/2* locus from anti-HA CUT&RUN for DMXAA-treated BMMs transduced with HA-SP140 or SP140, and ATAC-seq/RNA-seq of DMXAA-treated B6 and *Sp140^−/−^*BMMs. Alignments were visualized in the UCSC genome browser.

To test whether *Resist1/2* are direct targets of SP140, we performed CUT&RUN with an anti-HA antibody on DMXAA-treated *Sp140^−/−^*BMMs transduced with HA-SP140, or with untagged SP140 as a negative control. We confirmed that untagged and HA-tagged SP140 were functional and able to reduce late *Ifnb1* transcript levels induced by DMXAA (Extended Data Fig. 1a). In parallel, to characterize how SP140 regulates chromatin accessibility at target genes, we also generated ATAC-seq data from DMXAA-treated B6 and *Sp140^−/−^* BMMs. Consistent with previous SP140 ChIP-seq results^16^, we found that SP140 binds genetic loci involved in development, such as *Hoxa9* (Extended Data Fig. 1b-d). Similarly, we confirmed SP140 generally represses chromatin accessibility at target genes, consistent with previous studies implicating SP140 in transcriptional repression^15,16,18^ (Extended Data Fig. 1c). SP140-binding also correlated with the transcriptionally repressive histone mark H3K27me3 in publicly available ChIP-seq datasets (Extended Data Fig. 1e). However, a gene set previously reported to drive macrophage dysfunction after SP140 knockdown was not significantly upregulated in SP140-deficient BMMs^15,16^ (Fig. 2a, Extended Data Fig. 1f), suggesting that these genes do not explain elevated IFN-I induction that occurs in the absence of SP140. Consistent with our hypothesis that SP140 regulates *Ifnb1* mRNA stability and does not directly regulate *Ifnb1* transcription, SP140 did not bind or regulate chromatin accessibility at the *Ifnb1* gene (Extended Data Fig. 2a) or known *Ifnb1* regulatory elements^29–32^ (Extended Data Fig. 2b).

Importantly, SP140 robustly bound and repressed chromatin accessibility at the *Resist1/2* locus (Fig. 2b-c). Of the upregulated genes in DMXAA-treated *Sp140^−/−^Ifnar^−/−^* vs. *Ifnar^−/−^*BMMs, only *Resist1* was both bound by SP140 and showed increased chromatin accessibility in the absence of SP140 (Fig. 2b-c). SP140 binds a ∼10 kb region encompassing *Resist1* and *Resist2*, and negatively regulates chromatin accessibility at *Resist1/2* gene bodies (Fig. 2c). These results demonstrate that *Resist1* and *Resist2* are directly repressed by SP140 and are the only direct SP140 target genes upregulated along with *Ifnb1* in *Sp140^−/−^*macrophages.

### RESIST enhances *Ifnb1* mRNA stability

The RESIST protein has not been characterized in mice, but it is the ortholog of the human Annexin-2 Receptor (ANXA2R) which is encoded by a single *ANXA2R* gene. In wild-type mice, *Resist1* is detectably expressed in myeloid cells such as neutrophils, macrophages, monocytes, and eosinophils (Extended Data Fig. 3a). In human PBMCs, *ANXA2R* is detectably expressed in multiple immune cell populations, including monocytes, NK cells, DCs, ILCs, and T and B cells^33^ (Extended Data Fig. 3b). ANXA2R was first proposed as a receptor for Annexin-2 (ANXA2) based on the observation that ANXA2R overexpression increased the binding of exogenous ANXA2 to the cell surface^34^. However, ANXA2R is not likely to be a cell surface receptor as the DeepTMHMM algorithm^35^ fails to predict transmembrane helices. Moreover, recent work using unbiased mass spectrometry of immunoprecipitated human ANXA2R did not identify ANXA2 but instead identified the CCR4-NOT complex as the predominant ANXA2R-binding partner in cells^36^. We also found that a purified recombinant human ANXA2R protein did not robustly interact with the ANXA2-S100A complex, whereas a confirmed ANXA2 interaction partner (SMARCA3)^37^ was able to bind ANXA2-S100A (Extended Data Fig 3c).

CCR4-NOT is the major mRNA deadenylase complex in eukaryotic cells^38^. Specific mRNAs targeted for deadenylation are recruited to CCR4-NOT via mRNA-binding proteins (RBPs) that use small peptide motifs to dock to multiple interfaces on CCR4-NOT^38^. Recruited mRNAs are subsequently deadenylated, leading to their destabilization and degradation^38^. Human ANXA2R was proposed to inhibit CCR4-NOT and appeared to have antiviral effects^36^, though the antiviral mechanism was unclear.

As a unified hypothesis to explain how SP140 regulates *Ifnb1* mRNA stability and how ANXA2R provides antiviral defense, we propose that ANXA2R —which we now refer to as RESIST— stabilizes *Ifnb1* mRNA by binding CCR4-NOT to inhibit *Ifnb1* mRNA recruitment, deadenylation, and decay.

We first confirmed that HA-tagged mouse RESIST expressed in primary BMMs co-immunoprecipitated with CNOT1, the major scaffolding protein of CCR4-NOT (Fig. 3a). To test whether RESIST drives elevated *Ifnb1* mRNA levels in *Sp140^−/−^* BMMs, we disrupted *Resist1/2* genes by Cas9:gRNA electroporation of primary BMMs (Fig. 3b).

**Figure 3.**
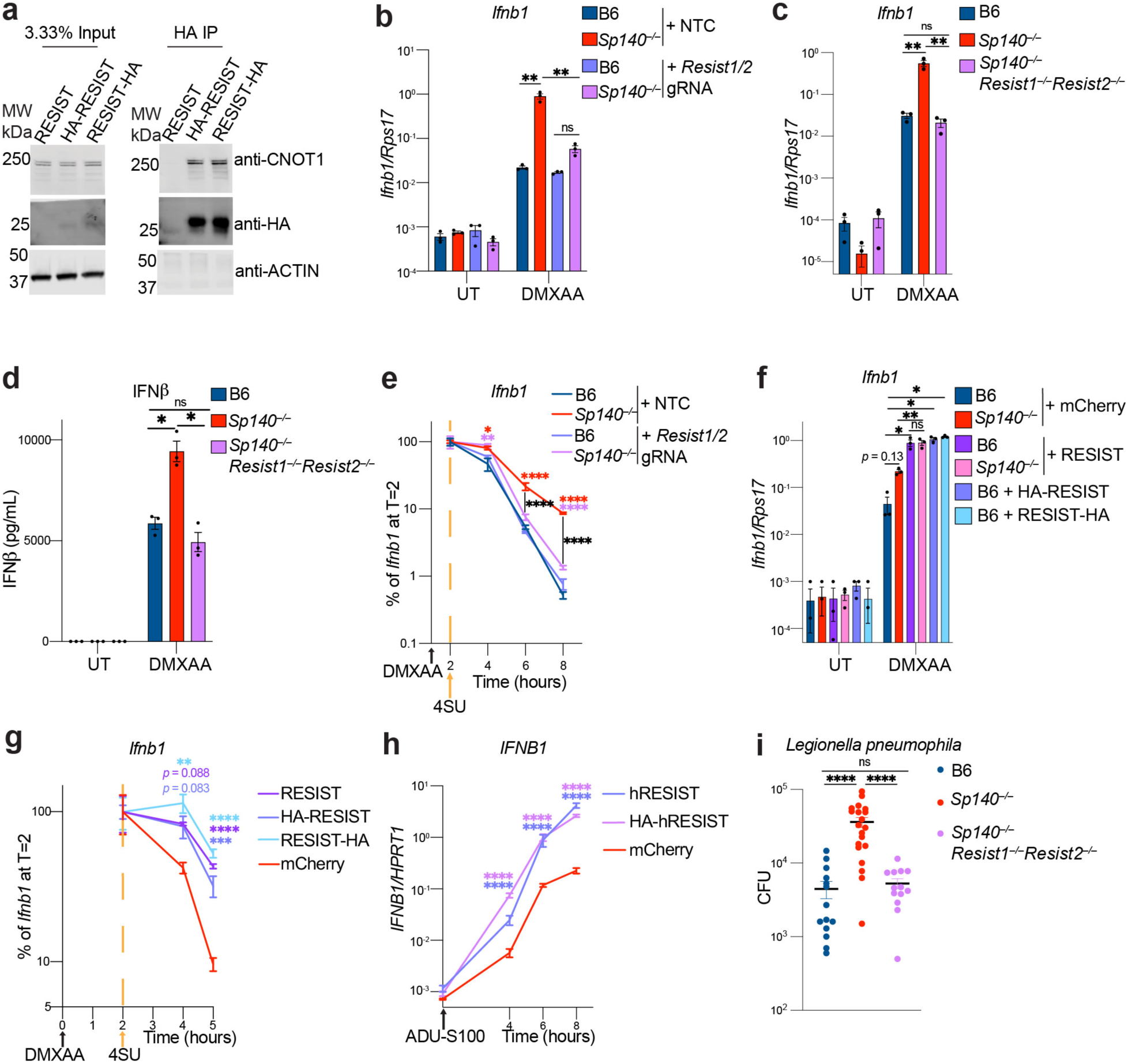
RESIST binds the CCR4-NOT complex and drives *Ifnb1* mRNA stabilization in *Sp140^−/−^* macrophages. a. Immunoblot of input lysate or anti-HA immunoprecipitate for BMMs transduced with lentiviral RESIST constructs, stimulated with doxycycline and 100 μg/mL DMXAA for 5-7 hours. For gel source data, please see Supplementary Figure 1. b. *Ifnb1* RT-qPCR from BMMs electroporated with non-targeting control (NTC) or *Resist1/2* gRNA, stimulated for 8 hours with 100 μg/mL DMXAA or untreated (UT). Knockout efficiency in the experiment shown was 86-87% for *Resist1* and 91% for *Resist2*. c. *Ifnb1* RT-qPCR from BMMs of indicated genotypes, stimulated for 8 hours with 100 μg/mL DMXAA or untreated (UT). d. ELISA of culture supernatant from BMMs treated with 100 μg/mL DMXAA for 24 hours. e. Roadblock RT-qPCR of BMMs electroporated with indicated gRNAs and treated with 100 μg/mL DMXAA, followed by 4SU at 2hr. The estimated knockout efficiency in the experiment shown was 71% for *Resist1* and 51-69% for *Resist2*. Asterisks indicate a significant difference in *Ifnb1* transcript compared to B6 + NTC at each time point and are colored by condition compared to B6 + NTC. Black bars/asterisks indicate a comparison between *Sp140^−/−^* + NTC and *Sp140^−/−^* + *Resist1/2* gRNA. f. *Ifnb1* RT-qPCR from BMMs transduced with indicated lentiviral constructs, treated with doxycycline and 100 μg/mL DMXAA for 7 hours. g. Roadblock RT-qPCR for *Ifnb1* from B6 BMMs transduced with indicated lentiviral constructs, stimulated with doxycycline and 100 μg/mL DMXAA at T0, followed by 4SU treatment at T2. Asterisks indicate a significant difference in *Ifnb1* compared to mCherry at each time point. h. *IFNB1* RT-qPCR from human BlaER1 monocytes overexpressing human RESIST constructs or mCherry, stimulated with ADU-S100 and doxycycline. Asterisks indicate a significant difference in *IFNB1* transcript compared to mCherry-expressing cells. i. Colony forming units (CFU) from lungs of mice infected with *Legionella pneumophila*, 96 hours post-infection. Results represent 3 independent pooled experiments, including an infection with *Sp140^−/−^Resist1^+/+^Resist2^+/+^ and Sp140^−/−^Resist1^−/−^Resist2^−/−^* littermates. * = *p* < 0.05, ** = *p* < 0.005, *** = *p* < 0.0005, **** = *p* < 0.00005, ns = not significant, calculated with one-way ANOVA tests with Dunnett’s T3 multiple comparison correction post-hoc (b, c, f) or FDR correction (d), or two-way ANOVA tests with Tukey’s post-hoc correction (e, g, h), or Mann-Whitney test (i).

*Resist1/2* disruption almost entirely eliminated the elevated *Ifnb1* transcript observed in DMXAA-treated *Sp140^−/−^* BMMs (Fig. 3b). We also generated *Sp140^−/−^ Resist1^−/−^ Resist2^−/−^*mice with CRISPR-Cas9 (Extended Data Fig. 3d), and confirmed that RESIST drove elevated *Ifnb1* transcript and IFNβ protein levels in *Sp140^−/−^* BMMs (Fig. 3c-d). Roadblock RT-qPCR confirmed that the decreased *Ifnb1* mRNA levels seen in *Resist1/2* knockout cells were due to decreased mRNA stability (Fig. 3e). Furthermore, RESIST overexpression in primary BMMs was sufficient to elevate *Ifnb1* transcript levels >10-fold, and eliminated the difference in *Ifnb1* transcript levels between B6 and *Sp140^−/−^*BMMs (Fig. 3f). RESIST overexpression also promoted *Ifnb1* transcript stabilization in primary BMMs, measured by Roadblock RT-qPCR (Fig. 3g).

Overexpressed human RESIST also drove >10-fold elevated *IFNB1* transcript levels in human BlaER1 monocytes, which do not endogenously express RESIST, upon stimulation with the STING agonist ADU-S100 (Fig. 3h). Finally, we found that *Resist1/2* entirely drove the susceptibility of *Sp140^−/−^* mice to *Legionella pneumophila* infection (Figure 3i), which we have previously shown to be largely IFN-I dependent^5^. These results identify mouse and human RESIST as a potent positive regulator of IFN-I.

### Mechanism of RESIST

To elucidate the mechanism by which RESIST stabilizes *Ifnb1* mRNA, we used AlphaFold-Multimer to generate predicted structures of the interaction between mouse RESIST and CCR4-NOT subunits implicated in IFN-I regulation, including 1) CNOT10/11, which recruit RBPs and negatively regulate IFN-I signaling in T-cells^39,40^, 2) the CNOT1 M-HEAT, which binds tristetraprolin (TTP), an RBP that has been implicated in negative regulation of *Ifnb1* mRNA stability^41–45^, and 3) CNOT9, which acts as a secondary binding site for TTP^46^ and was shown to directly bind ROQUIN1, which, along with ROQUIN2, negatively regulates transcript stability for pro-inflammatory cytokines^11,47^. Our predicted structures suggested that RESIST may interact with multiple sites on the CCR4-NOT complex (Fig. 4a, Extended Data Fig. 4a). Such multivalent interactions are typical of many CCR4-NOT interacting proteins^38^. Intriguingly, the RESIST C-terminal region is predicted to fold into a helix (residues 168-177) and to interact with CNOT1 M-HEAT at the site previously shown to be bound by TTP^41^ (Fig. 4a, Extended Data Fig. 4). Moreover, RESIST was predicted to bind the tryptophan binding pockets on CNOT9 also known to interact with TTP (Fig. 4a, Extended Data Fig. 5)^46^. Both interactions were characterized by hydrophobic interactions between RESIST and CNOT1/CNOT9, as well as a high number of predicted models and predicted local distance difference test (pLDDT), and low predicted aligned error (PAE) (Extended Data Fig. 4b-e and 5a-d). AlphaFold also predicted, with lower confidence, that the RESIST C-terminal region interacted with a CNOT9 interface bound by *Drosophila* Roquin^47^ (Fig. 4a, Extended Data Fig. 5a-d). In contrast, AlphaFold did not predict a high-confidence interaction between RESIST and CNOT10/11 (Extended Data Fig. 6).

**Figure 4.**
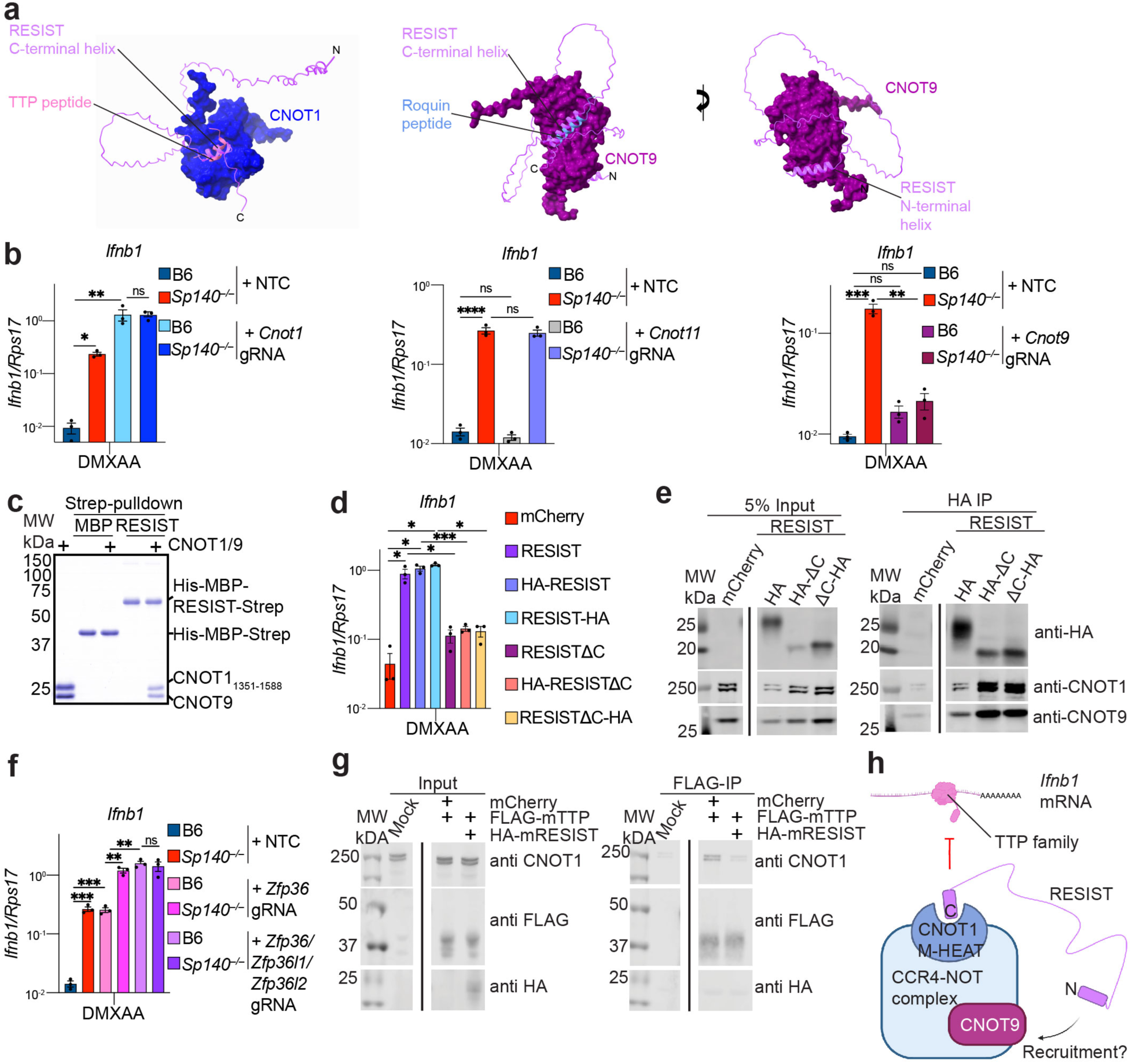
RESIST counteracts repression of IFN-I by TTP family proteins, a function which requires a RESIST C-terminal region and CNOT9. a. AlphaFold predictions of RESIST with CNOT1 M-HEAT and CNOT9. TTP peptide:CNOT1 M-HEAT is PDB code 4J8S^41^. Roquin peptide:CNOT9 is PDB code 5LSW^47^. N and C-terminal ends of RESIST are marked by “N” or “C” respectively. b. *Ifnb1* RT-qPCR from BMMs electroporated with non-targeting control gRNA (NTC) or gRNAs targeting indicated CCR4-NOT subunits, treated for 8 hours with 100 μg/mL DMXAA. c. Strep pulldown of purified recombinant full-length human His-MBP-RESIST-Strep or His-MBP-Strep with human CNOT9 and CNOT1 (aa 1351-1588). First lane indicates purified CNOT9/CNOT1. d. *Ifnb1* RT-qPCR for BMMs transduced with indicated lentiviral constructs, treated with doxycycline and DMXAA for 6 hours. Results include data also shown in Fig. 3f. e. Immunoblot for anti-HA IP of BMMs transduced with indicated constructs in d. RESIST construct is C-terminally tagged with HA. For gel source data, please see Supplementary Figure 1. f. *Ifnb1* RT-qPCR from BMMs electroporated with indicated gRNAs, treated for 8 hours with 100 μg/mL DMXAA. g. Immunoblot for FLAG IP of FLAG-TTP (mouse) co-expressed with mouse HA-RESIST in HEK293Ts. For gel source data, please see Supplementary Figure 1. h. Schematic for how RESIST may interact with CCR4-NOT subunits CNOT1 and CNOT9 to mediate stabilization of *Ifnb1* mRNAs. * = *p* < 0.05, ** = *p* < 0.005, *** = *p* < 0.0005, **** = *p* < 0.00005, ns = not significant, calculated with one-way ANOVA tests with Dunnett’s T3 multiple comparison correction post-hoc.

To assess the functionality of CCR4-NOT subunits predicted to interact with RESIST, we genetically disrupted CCR4-NOT subunits (*Cnot1/11/9*) in BMMs with Cas9:gRNA electroporation. We evaluated *Ifnb1* transcript levels by RT-qPCR at the late 8 hr time point after DMXAA stimulation (Fig. 4b). Notably, targeting of the CCR4-NOT scaffold gene *Cnot1* was not well-tolerated, with 5-10-fold fewer cells recovered after electroporation relative to non-targeting controls (NTC), and variable knockout efficiency between experiments. Nevertheless, *Cnot1* deficiency phenocopied RESIST overexpression and led to similarly increased *Ifnb1* upon DMXAA stimulation in both B6 and *Sp140^−/−^* cells (Fig. 4b, Extended Data Fig. 7a), consistent with the hypothesis that RESIST elevates *Ifnb1* transcript levels by inhibition of CCR4-NOT. By contrast, targeting *Cnot11* did not regulate *Ifnb1* in either B6 or *Sp140^−/−^* BMMs (Fig. 4b, Extended Data Fig. 7a), demonstrating that RESIST-mediated *Ifnb1* mRNA stability does not require CNOT11, and consistent with the absence of a predicted interaction.

Interestingly, deletion of *Cnot9* eliminated elevated *Ifnb1* transcript in DMXAA-stimulated *Sp140^−/−^* BMMs, but did not affect *Ifnb1* levels in DMXAA-stimulated B6 BMMs (Fig. 4b, Extended Data Fig. 7a). These results indicate CNOT9 is required for RESIST function. Using purified recombinant RESIST and CNOT9 proteins, we were able to observe a direct stoichiometric interaction between RESIST and CNOT9 (Fig. 4c).

To probe the functional significance of the RESIST C-terminal region, we generated a truncation mutant (RESISTΔC). BMMs transduced with the RESISTΔC mutant induced much lower levels of *Ifnb1* transcript than BMMs transduced with WT RESIST (Fig. 4d), suggesting the C-terminal region is essential for RESIST-mediated *Ifnb1* mRNA stabilization. Interestingly, the RESISTΔC mutant still immunoprecipitated with the CCR4-NOT complex (Fig. 4e), consistent with a predicted multivalent interaction between RESIST and the CCR4-NOT complex (Fig. 4a).

RESIST is predicted to interact with the known interfaces that TTP and Roquin bind on the CCR4-NOT complex to mediate the decay of their target transcripts. We therefore asked whether TTP and/or Roquin mediated *Ifnb1* mRNA decay. We disrupted the genes encoding Roquin1/2 (*Rc3h1/2)*, as well as TTP (*Zfp36*) alone and in combination with additional TTP family members (*Zfp36l1, Zfp36l2)*, and evaluated *Ifnb1* transcript levels by RT-qPCR. Genetic disruption of *Rc3h1/2* did not affect *Ifnb1* induced by DMXAA in either B6 or *Sp140^−/−^*BMMs (Extended Data Fig. 7c-d). By contrast, loss of *Zfp36* in DMXAA-stimulated B6 BMMs increased *Ifnb1* transcript levels (Fig. 4f, Extended Data Fig. 7b). We validated that the RNA-binding zinc finger domain of TTP was able to specifically bind the *Ifnb1* 3’UTR in a manner dependent upon an AU-rich element (ARE) that is a canonical TTP target sequence (Extended Data Fig. 8). However, *Sp140* deficiency further increases *Ifnb1* in the absence of *Zfp36*, implying that RESIST has additional activities beyond TTP inhibition in *Sp140^−/−^* BMMs (Fig. 4f, Extended Data Fig. 7b). Consistent with this hypothesis, deletion of the genes encoding the additional TTP family members *Zfp36l1/Zfp36l2* resulted in further elevation of *Ifnb1* (Fig. 4f, Extended Data Fig. 7b). These results imply a previously unappreciated role for these TTP family members in negative regulation of *Ifnb1* mRNA stability. Crucially, loss of *Sp140* does not affect *Ifnb1* levels in the absence of the *Zfp36* gene family (Fig. 4f).

These results are consistent with a model in which RESIST interferes with the ability of TTP family members to mediate *Ifnb1* transcript destabilization. Indeed, we found RESIST inhibited the interaction of TTP with CNOT1 when both RESIST and TTP were co-expressed in 293T cells (Fig. 4g). A potential model consistent with our data is one in which RESIST binds CCR4-NOT and hinders the interaction between the CCR4-NOT complex and TTP family members, leading to *Ifnb1* mRNA stabilization (Fig. 4h). In contrast, CNOT9, which is required for elevated *Ifnb1* in *Sp140^−/−^* cells, appears to promote RESIST function, potentially by facilitating the interaction of RESIST with CCR4-NOT (Fig. 4h). A detailed mechanism for RESIST function will require structural analyses of the RESIST-CCR4-NOT complex.

### Antiviral activity of SP140

Lastly, we considered why SP140 may have evolved to repress the antiviral cytokine IFN-I. Intriguingly, SP140 is part of a family of proteins that includes SP100, a well-described antiviral protein^48^, and SP140L, which was recently described to have an antiviral role against the herpesvirus Epstein-Barr virus^49^. Mechanistically, SP100 is thought to repress transcription of viral genomes in the nucleus through sequestration in transcriptionally repressive nuclear bodies (NBs) that co-localize with the NB protein PML^48^. Whether SP140 co-localizes with PML or has antiviral activity is unclear^4,50-53^.

Importantly, because SP100 and other NBs are often antiviral, viruses often encode effectors to disrupt NB function^29,48,54^. To counter the effector-mediated disruption of NBs, we previously proposed that antiviral NB proteins can evolve the ability to repress IFN-I as a secondary function^6,29^. Thus, viruses that disrupt NBs to evade their primary antiviral activity will unleash a secondary “backup” interferon response. The secondary response is sometimes called “effector-triggered immunity”, a major immune strategy in plants^7^ that remains poorly characterized in mammals^6^.

To test the hypothesis that SP140 is an antiviral nuclear body protein, we first characterized the nature of SP140 NBs. To detect endogenously expressed SP140, we generated *HA-Sp140* knock-in mice by inserting an HA-tag after the start codon of the *Sp140* gene (Extended Data Fig. 9a). Immunoblot of BMMs from *HA-Sp140^+/+^* mice confirmed the presence of an anti-HA reactive band at the expected molecular weight of SP140 (Extended Data Fig. 9b). HA-SP140 was expressed at similar levels as WT SP140, and was functional, as assessed by its ability to repress *Ifnb1* at late timepoints in DMXAA-treated BMMs (Extended Data Fig. 9b-c). Surprisingly, immunofluorescence of *HA-Sp140^+/+^* BMMs demonstrated that SP140 forms large NBs that do not co-localize with PML bodies (Fig. 5a, Extended Data Fig. 9d). Rather, we found that SP140 NBs appeared to largely overlap with the nucleolar marker fibrillarin (Fig. 5b, Extended Data Fig. 9e).

**Figure 5.**
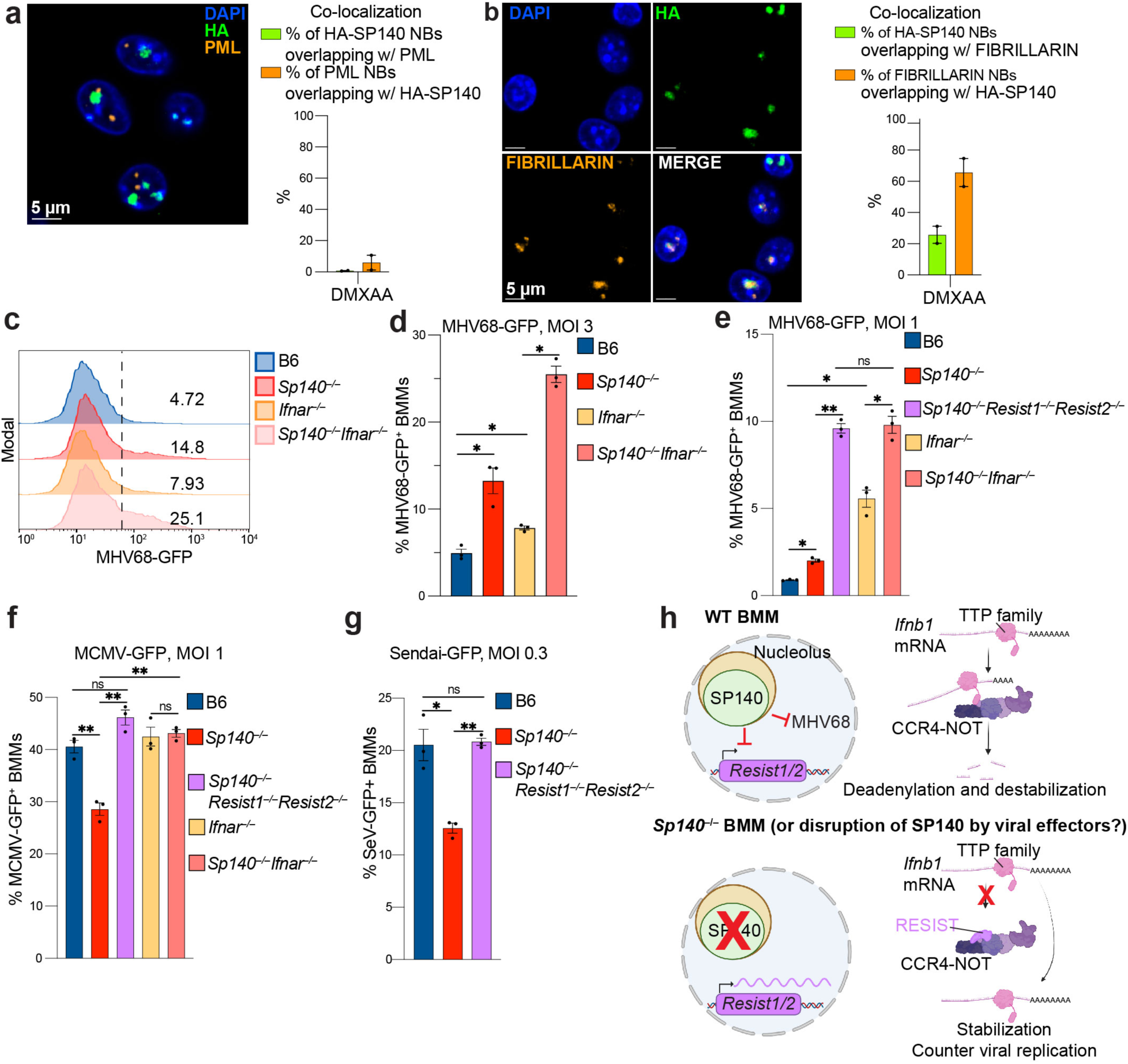
SP140 is an antiviral NB protein that co-localizes with nucleoli. a. Left: Immunofluorescence of *HA-Sp140^+/+^* BMMs treated with DMXAA and stained with DAPI, anti-HA, and anti-PML. Right: quantification of overlap of HA-SP140 and PML NBs for 2 independent experiments. b. Immunofluorescence of *HA-Sp140^+/+^* BMMs treated with DMXAA and stained with DAPI, anti-HA, and anti-fibrillarin. Right: quantification of overlap of HA-SP140 and fibrillarin NBs for 2 independent experiments. c. Histograms of MHV68-GFP signal in MHV68-GFP infected BMMs, assessed by flow cytometry. Numbers represent MHV68-GFP^+^ as a percentage of live cells. d. Quantification of MHV68-GFP^+^ cells from c). e. Quantification of MHV68-GFP+ BMMs. f. Quantification of MCMV-GFP+ BMMS, assessed by flow cytometry. g. Quantification of Sendai-GFP+ BMMs, assessed by flow cytometry. h. Schematic for the proposed model of SP140 antiviral activity and RESIST-mediated stabilization of *Ifnb1* transcript. * = *p* < 0.05, ** = *p* < 0.005, *** = *p* < 0.0005, **** = *p* < 0.00005, ns = not significant, calculated with one-way ANOVA tests with FDR correction (d-g).

We then tested whether SP140 has antiviral activity. We infected B6 and *Sp140*^−/−^ BMMs with MHV68-GFP, a mouse herpesvirus engineered to encode GFP. We found that a larger fraction of *Sp140*^−/−^ BMMs were GFP^+^ compared to WT B6 BMMs (Fig. 5c-d, Extended Data Fig. 10a). The antiviral effect of SP140 was independent of IFN-I signaling, as *Sp140^−/−^Ifnar^−/−^* BMMs also exhibited higher levels of infection as compared to *Ifnar^−/−^* BMMs (Fig. 5c-d). We also observed that *Sp140^−/−^* BMMs exhibited higher *Ifnb1* transcript levels than wildtype cells upon MHV68-GFP infection (Extended Data Fig. 10b). Consistent with this observation and the role of RESIST as a critical driver of IFN-I in *Sp140*^−/−^ BMMs, *Sp140^−/−^Resist1^−/−^Resist2^−/−^* BMMs were more susceptible to MHV68-GFP infection than *Sp140^−/−^* BMMs, and phenocopied the susceptibility of *Sp140^−/−^Ifnar^−/−^* BMMs (Fig. 5e). These data suggest that elevated, RESIST-dependent IFN-I in *Sp140*^−/−^ BMMs counteracts the loss of the antiviral protein SP140 during MHV68-GFP infection.

We then tested whether SP140 has antiviral activity against other viruses. We infected B6 and *Sp140*^−/−^ BMMs with MCMV-GFP, another mouse herpesvirus engineered to encode GFP. In contrast to MHV68, we found that MCMV-GFP was modestly restricted in *Sp140*^−/−^ BMMs (Fig. 5f). The restriction of MCMV-GFP in *Sp140*^−/−^ cells depended on RESIST and IFNAR (Fig. 5f). We also found that *Sp140*^−/−^ BMMs restricted the replication of an RNA-virus, Sendai virus, also encoding GFP (Fig. 5g), whereas *Sp140*^−/−^*Resist1^−/−^Resist2^−/−^* BMMs and WT BMMs exhibited similar levels of Sendai-GFP replication. These results suggest the antiviral activity of SP140 is specific to certain viruses such as MHV68-GFP, and that the elevated IFN-I due to RESIST expression in the absence of SP140 counters replication of diverse viruses.

## Discussion

Our results describe several new findings. First, we identify RESIST as a novel positive regulator of type I IFN, which we find operates via the previously undescribed mechanism of promoting *Ifnb1* mRNA stability. We propose that RESIST may represent a founding example of a cytokine regulator that functions via CCR4-NOT inhibition.

Second, we demonstrate that RESIST is a direct target of SP140 transcriptional repression, and that RESIST drives the enhanced *Ifnb1* levels in *Sp140*^−/−^ cells. Third, we demonstrate that the RNA binding protein TTP (ZFP36), and its previously poorly described paralogs ZFP36L1 and ZFP36L2, are important but partly redundant negative regulators of IFN-I. Lastly, we describe an antiviral function for SP140 against MHV68 that is independent of its ability to regulate IFN-I.

At present, the mechanism by which SP140 is antiviral is unknown. SP140 contains domains homologous to those found in the antiviral protein SP100. Like SP100, the DNA-binding SAND domain of SP140 may also bind viral genomes, while the CARD domain may mediate SP140 oligomerization into transcriptionally repressive nuclear bodies^48^. Interestingly, our results suggest SP140 co-localizes distinctly from SP100/PML, suggesting the antiviral mechanism of SP140 may be distinct from that of SP100. How SP140 is antiviral will be of great interest to determine in future work.

Our data support a model whereby RESIST counteracts destabilization of *Ifnb1* mRNA mediated by CCR4-NOT and the TTP-family. The activity of RESIST requires its C-terminal region. We predict that RESIST interacts through multiple binding sites with the CCR4-NOT complex. Interestingly, the predicted sites of RESIST interaction overlap with known regions of interaction between CCR4-NOT and TTP. Validation of the predicted interactions between RESIST and CCR4-NOT, and a detailed investigation of whether RESIST competes with TTP family members for CCR4-NOT interaction, will require further work including the structural analyses of the RESIST-CCR4-NOT complex.

Our data do not exclude a model in which RESIST stabilizes all CCR4-NOT mRNA targets by acting as a non-specific inhibitor of deadenylation, similar to RNF219^55^. However, in *Sp140*-deficient cells, we observe few upregulated transcripts other than *Resist1* and *Ifnb1* (Fig. 2a), suggesting that RESIST acts to stabilize a specific set of targets. Curiously, we also found that TTP-regulated transcripts like *Tnf* and *Il6* are not derepressed in *Sp140^−/−^* macrophages (Fig. 2a). The apparent specificity of RESIST could derive from multiple sources, including cellular co-localization of RESIST and *Ifnb1* mRNA transcripts, preferential targeting of *Ifnb1* mRNA by the TTP family members, or particular sensitivity of *Ifnb1* transcripts to the partial TTP inhibition mediated by RESIST. It is also possible that RESIST may regulate other transcripts besides *Ifnb1 in vivo*. Ultimately, further work is required to determine the mechanistic basis for the apparent specificity of RESIST for *Ifnb1* in macrophages, and whether RESIST regulates other transcripts *in vivo*.

While this work focuses on the SP140-RESIST circuit, RESIST may also function independently of SP140. In human cells, RESIST is induced by IFN-I^36^, which suggests that it may serve an antiviral function. It remains unknown if human SP140 or other SP family members also regulate human *RESIST.* In mice, we do not observe destabilization of *Ifnb1* transcripts in *Resist1/2^−/−^* BMMs that are *Sp140*^+^, likely because SP140 strongly represses RESIST expression in WT macrophages. However, it is possible that RESIST is regulated differently in other cell types *in vivo*.

Our results highlight an important role for TTP and TTP family members ZFP36L1/ZFP36L2 in IFN-I repression in macrophages. Our finding that TTP represses IFN-I is consistent with previous work that suggests TTP negatively regulates the stability of *Ifnb1* mRNA^42,43,45,56^. TTP detectably binds the 3’UTR of *Ifnb1* mRNA (Extended Data Fig. 8)^42^, and a hyperactive TTP mutant represses *Ifnb1* transcript levels in mouse macrophages^43^. ZFP36L1/ZFP36L2 are understudied, and genetic deletion in mice is either embryonic or perinatally lethal^44^. However, ZFP36L1/ZFP36L2 are known to recruit CCR4-NOT to target mRNAs and modulate B/T cell function^44,57-59^. Interestingly, while *Zfp36^−/−^* mice develop severe autoinflammation, mice lacking myeloid-specific TTP expression do not develop immunopathology^44^, suggesting functional redundancy among TTP family members in myeloid cells. Indeed, recent work described fatal autoinflammation in mice lacking myeloid-specific expression of *Zfp36/Zfp36l1/Zfp36l2*, which may result partly from elevated *Ifnb1* transcripts in these mice^56^.

Our results demonstrate that SP140 largely resides in nucleoli and is intrinsically antiviral against MHV68 (Fig. 5d-e). In addition, we show that SP140 has evolved a secondary function: repression of the *Resist1/2* genes, which encode a previously undescribed positive regulator of IFN-I. We propose that these apparently conflicting anti– and pro-viral roles of SP140 can be explained by a host-virus evolutionary arms race in the nucleus (Fig. 5h). Many viral effectors target antiviral nuclear bodies^29,48,54^. We speculate that SP140 evolved to protect itself from attack by acquiring the ability to negatively regulate RESIST. In this scenario, viral attack of SP140 would de-repress RESIST, which would augment IFN-I mediated antiviral defense. Thus, the SP140-RESIST pathway may provide an example of effector-triggered immunity^6,7,29^.

Finally, we speculate that the antiviral activity of SP140 could explain why multiple sclerosis and B-cell cancers are linked to LOF *SP140* mutations, as infection with MHV68-related viruses Epstein-Barr Virus (EBV) and Kaposi’s sarcoma-associated herpesvirus (KSHV) are associated with these immune disorders^60–62^. In future work, it will be of interest to determine whether individuals with LOF *SP140* mutations are more susceptible to EBV/KSHV infection, which could lead to increased risk of MS and B-cell cancers.

## Methods

### Mice

All animal experiments complied with the regulatory standards of, and were approved by, the University of California Berkeley Institutional Animal Care and Use Committee. Mice were maintained in specific-pathogen-free conditions with a 12-hour light/dark cycle and given water and standard chow diet (Harlan’s irradiated chow) *ad libitum*. All mice were bred in-house. C57BL/6 and B6.129S2-*Ifnar1^tm1Agt^*/Mmjax (*Ifnar^−/−^)* were originally from Jackson laboratories and MMRC and further bred in-house. The generation and genotyping of *Sp140^−/−^* mice was previously described^5^. *HA-Sp140* knock-in mice were generated by electroporation of C57BL/6 zygotes with Cas9, Alt-R® CRISPR-Cas9 crRNA from IDT (gRNA sequence: ACUCCAAGGGACCCUGUUCA), and a homology repair Alt-R HDR donor oligo from IDT (CCCCTGAAGGAGTTCTCTCTGGGCTTCCCAGAGACTCAGAGGGGGTTCGGTCTAGTCTGAACAGGGTCCCTTGGAGTCTGTGTAGGGGATGTACCCATACGATGTTCCAGATTACGCTGCAGGAGGCTACAATGAACTCAGCAGCAGGTAAGTCCCATTCTCTCTTGTCCCTTGTCTC) as previously reported^63^. Founders were backcrossed to C57BL/6J, and mice with matching *HA-Sp140* alleles were further bred. *HA-Sp140* knock-in mice were genotyped using a qPCR-based assay from Transnetyx. *Sp140^−/−^ Resist1^−/−^Resist2^−/−^* mice were generated by electroporation of *Sp140^−/−^* zygotes with Cas9 and sgRNA (CGACGATGGCGGTGACTACC)^5^. Founders were genotyped and backcrossed to *Sp140^−/−^* mice, and progeny with matching alleles were further bred. For genotyping *Resist1/*2, large ear clips were obtained and digested in QuickExtract lysis buffer overnight with 0.4 mg/mL Proteinase K, followed heat inactivation (85°C for 4 min, 98°C for 2 min) and stored at –80. After freezing, ear clip lysates were vortexed. Mice were genotyped for *Resist1* (*Gm21188*) with Q5 2X PCR mix according to manufacturer instructions with 1 μL of lysate per 20 μL PCR reaction (F: TTGAGAAATCCGTTTGTAATGGG, R: GCCTTTCTCCGGATTCACGA; cycling conditions: 98°C for 3 min, 35 cycles of 98°C for 10 s, 63.5°C for 20 s, 72°C for 65s, followed by a final extension of 72°C for 10 min). Mice were genotyped for *Resist2* (*Gm36079*) with Phusion GC rich PCR components according to manufacturer instructions, using 1 μL of lysate per 20 μL PCR reaction (F: TGGTATTCTCTAGAGATAACATCACAGCACCTACTTACTCC, R: CCTCCCCTCGCCATCACTGCCTG; cycling conditions: 98°C for 30s, 30 cycles of 98°C for 10 s, 72°C for 15 s, 72°C for 60s, followed by a final extension of 72°C for 10 min). Five μL of PCR product was cleaned with FastAP and ExoI, then diluted 2-fold with water and Sanger sequenced *(Resist1* R: GCCTTTCTCCGGATTCACGA; *Resist2* F seq: CTGAATGATTCTTCTACTGCTTCCATCC).

### Cell culture

HEK293T and GP2 cells were obtained from the UC Berkeley Tissue Culture facility and further propagated at 37°C and 5% CO_2_ in complete DMEM (10% FBS v/v (Gibco) and 1X pen-strep and glutamine (Gibco)). BlaER1 cells in B cell stage were cultured in complete RPMI, and differentiated into monocytes as described^29^. Cell lines were tested by PCR for mycoplasma (F: CACCATCTGTCACTCTGTTAACC, R: GGAGCAAACAGGATTAGATACCC) with Dreamtaq PCR reagents and validated with STR profiling by the UC Berkeley Tissue culture facility.

### BMM generation and stimulation

Mice were euthanized and bones (femurs and tibias) were extracted with stringent washing in 70% ethanol. After washing in 70% ethanol and BMM media (complete RPMI with 10% MCSF (v/v) generated from 3T3 cells as described^5^), bones were crushed in a sterilized mortar and pestle and bone marrow was passed through a 70 μm filter. Bone marrow from one mouse was divided across 8 15 cm non-TC-treated plates in 30 mL total volume in BMM media. Day of bone marrow harvest was considered as day 0, and BMMs were fed on day 3 with 10 mL media per plate. On day 6, BMMs were harvested in cold PBS by scraping and seeded on non-TC treated plates at the appropriate density (100,000 cells per non-TC 12-well and 24-well; 6-well for over 100,000 cells/well) and rested at least overnight before stimulation. To stimulate, media was aspirated and replaced with BMM containing 10 ng/mL LPS (Invivogen, tlrl-3pelps), 100 μg/mL poly(I:C) (Invivogen, tlrl-picw), or 100 μg/mL DMXAA (Cayman Chemicals, 14617).

### RT-qPCR

RNA was isolated with the Omega Biotek Total RNA II kit according to kit instructions. RNA was subsequently DNase-treated either on-column (Qiagen, 79254) or with RQ1 (Promega, M6101) according to manufacturer instructions. For BlaER1 cells, RNase-inhibitor was always included with DNase treatment (either RNaseOUT (Invitrogen) or RNasin (Promega)). RNA was then converted to cDNA with Superscript III Reverse Transcriptase (Invitrogen, 18080093) and oligo dt_18_ (NEB, S1316S) in the presence of RNase-inhibitors. Diluted cDNA was assessed by qPCR with Power SYBR™ Green PCR Master Mix (Fisher Scientific, 43-676-59) reagents, using technical duplicates. A standard curve generated from samples within each experiment was used to quantify relative amounts of transcript. *Ifnb1* transcript levels (F: GTCCTCAACTGCTCTCCACT, R: CCTGCAACCACCACTCATTC) were normalized to housekeeping genes including *Rps17* (F: CGCCATTATCCCCAGCAAG, R: TGT CGGGATCCACCTCAATG), *Oaz1* (F: GTGGTGGCCTCTACATCGAG, R: AGCAGATGAAAACGTGGTCAG), or *Hprt1* (F: GTTGGATACAGGCCAGACTTTGTTG, R: GAGGGTAGGCTGGCCTATAGGCT) as indicated in figures. Human *IFNB1* transcript quantities (F: CAGCATCTGCTGGTTGAAGA, R: CATTACCTGAAGGCCAAGGA) were normalized to *HPRT1* (F: ATCAGACTGAAGAGCTATTGTAATGA, R: TGGCTTATATCCAACACTTCGTG). DNase-treated RNA was also included in qPCR assays to ensure complete digestion of genomic DNA. Wells with a poor ROX reference were excluded from analysis, as were samples with unusually low housekeeping gene amounts indicative of RNA degradation. Replicates in figures indicate biological replicates (separate wells).

### Roadblock RT-qPCR

BMMs were either pre-treated with freshly prepared 400 μM 4-thiouridine (4SU) (Cayman Chemicals, 16373) 1-2 hours before 100 μg/mL DMXAA stimulation or treated with 4SU 2 hours after DMXAA stimulation, as previously described^28,64^. Briefly, RNA was harvested as described above and DNase-treated. Equal amounts of RNA (∼200 ng or more) was treated with 48 μM N-ethyl maleimide (Sigma, Cat. No. 04259-5G), and quenched with 20 mM DTT. RNA was then purified with RNAClean beads (Beckman Coulter, A63987) and converted to cDNA with ProtoScript® II Reverse Transcriptase according to manufacturer instructions in the presence of RNAse inhibitors. Quantitative PCR was then performed as described above, with primers for *Ifnb1* (F: TGGATGGCAAAGGCAGTGTAA, R: CACCTACAGGGCGGACTTC) and *Rps17* (see above). In some experiments, house-keeping normalized *Ifnb1* is displayed in figures as the percentage of average *Ifnb1/Rps17* quantity present at 2 hours of DMXAA stimulation for each condition. To validate the Roadblock RT-qPCR protocol, in every Roadblock RT-qPCR experiment, BMMS were also pre-treated in parallel with 4SU before DMXAA treatment, and RNA was isolated at T=2 and other timepoints indicated in figures and converted to cDNA. NEM treatment was confirmed to reduce detection of *Ifnb1* transcript by qPCR ∼10-fold with 4SU pre-treatment before DMXAA for every Roadblock RT-qPCR experiment.

### ELISA

BMMs were seeded at 85,000 cells per well in non-TC treated 96-well plates in 200 μL of medium and rested overnight. Cells were stimulated with indicated stimuli in figure legends for 24 hours. Plates were spun at 600 *x* g for 5 minutes, and supernatants were removed and stored at –80. Culture supernatants from DMXAA-treated cells were diluted 1:50-1:100 before evaluation with the Lumikine Xpress mIFN-β 2.0 kit according to manufacturer instructions.

### RNA-seq sample generation and analysis

Day 6 BMMs derived from B6 and *Sp140^−/−^,* or *Ifnar^−/−^* and *Sp140^−/−^Ifnar^−/−^*mice were seeded at 1 e6/well in 6 well non-TC plates and rested overnight. BMMs were then either left unstimulated or stimulated with DMXAA (100 μg/mL for B6 and *Sp140^−/−^*BMMs, and 10 μg/mL DMXAA for *Ifnar^−/−^* and *Sp140^−/−^Ifnar^−/−^* BMMs) for either 8 hours for B6 and *Sp140^−/−^* BMMs, or 4 hours for *Ifnar^−/−^* and *Sp140^−/−^Ifnar^−/−^* BMMs. RNA was isolated with the Omega Biotek Total RNA II kit, and DNAse-treated with TURBO DNAse (ThermoFisher Scientific, AM2238) according to manufacturer instructions. For RNA from B6 and *Sp140^−/−^*BMMs, single-index libraries were generated by Azenta, using rRNA depletion (Qiagen, QIAseq FastSelect – HMR rRNA Removal kit) and the NEBNext RNA Ultra kit (NEB). Libraries were evenly split across 2 lanes of an Illumina Hiseq flow cell and sequenced (150 bp paired-end reads, depth of 25-30M reads/sample). For *Ifnar^−/−^* and *Sp140^−/−^Ifnar^−/−^* BMM samples, libraries were prepared for Illumina sequencing with dual indices using polyA selection and KAPA HyperPrep reagents (Roche) after heat fragmentation through the UC Berkeley QB3 Vincent J. Coates Genomics Sequencing Laboratory. Libraries were subsequently size selected between 450-500 bp, and then sequenced on an Illumina Novaseq flow cell for a depth of over 20M mapped reads (150 bp, paired-end). B6 and *Sp140^−/−^* samples were collected separately from *Ifnar^−/−^* and *Sp140^−/−^Ifnar^−/−^*samples. Three biological replicates for each genotype and condition were collected across three independent experiments from 3 separate mice for each genotype. Adapters and low-quality reads were trimmed using BBDuk v38.05 with arguments ‘ktrim=r k=23 mink=11 hdist=1 mapq=10 qtrim=r trimq=10 tpe tbo’. For UCSC genome browser visualization, reads were mapped to mm10 using hisat2 v2.1.0 with options ‘--no-softclip –k 100 | samtools view –q 10 –Sb – | samtools sort’. CPM normalized bigwigs were made using deepTools bamCoverage v3.0.1. For transcript quantification, reads were mapped to mm10 (gencode.vM18.annotation.gtf) using using Salmon v0.13.1 with options ‘--libType A –-validateMappings –-rangeFactorizationBins 4 –-gcBias’. Differentially expressed genes were called using DESeq2 v1.38.3 with design ‘∼batch + genotype + treatment + genotype:treatment’. For differential expression analysis of *Sp140^−/−^* vs. B6 BMMs and *Sp140^−/−^Ifnar^−/−^* vs. *Ifnar^−/−^* BMMs, normalized count data was derived from the following DESeq2 comparisons: 1) SP140-deficient BMMs treated with DMXAA (3 replicates) vs. SP140-WT BMMs treated with DMXAA (3 replicates); and 2) untreated SP140-deficient BMMs (3 replicates) vs. untreated SP140-WT BMMs (3 replicates). Genes with zero counts across all samples were removed. Volcano plots were generated with ggplot2 v3.5.0 R package.

### ATAC-seq sample generation and analysis

B6 and *Sp140^−/−^* samples BMMs were derived and stimulated as described above with 100 μg/mL DMXAA for 8 hours in three separate experiments, in parallel to samples generated for RNA-seq. BMMs were harvested in PBS and counted, and ATAC-seq samples were generated from 100,000 input cells essentially as described^65^, except that isolated nuclei were spun at 1000 *x g*. Illumina-compatible libraries were prepared as described^65^, with additional Ampure XP bead (Beckman Coulter) purification to remove contaminating adaptor dimers. Samples were sequenced with Azenta on an Illumina HiSeq flow cell (over 50M paired end reads/sample, 150 bp reads). Adapters and low quality reads were trimmed using BBDuk v38.05 with arguments ‘ktrim=r k=23 mink=11 hdist=1 maq=10 qtrim=r trimq=10 tpe tbo’ and mapped to mm10 using BWA-MEM v0.7.15, and only uniquely mapping reads with a minimum MAPQ of 10 were retained.

Fragments aligning to the mitochondrial genome were removed. Peak calling was performed using complete and size subsetted alignment files with MACS2 v2.1.1 with paired-end options ‘--format BAMPE –-SPMR –B –-broad’. For visualization, CPM normalized bigwigs were made using deepTools bamCoverage v3.0.1.

### BMM transduction

Low passage HEK293T cells or GP2s were seeded in a 6-well format on tissue culture treated plates and rested at least overnight. To generate lentivirus, HEK293Ts at >70% confluency in a 6-well TC-treated plate were transfected with 0.468 μg VSV-G (pMD2.G, Addgene plasmid #12259), 1.17 μg D8.9 packaging vector, and 1.56 μg of doxycycline-inducible, puromycin-selectable lentiviral vector per well using Lipofectamine 2000 according to manufacturer instructions. The lentiviral backbone used in this study (pLIP) was adapted from pLIX (Addgene #41394) by removal of ccdB^29^. Gene blocks encoding codon-optimized mouse/human RESIST, or mCherry were cloned into pLIP after the dox-inducible promoter, digested with NheI and BamHI, with Infusion reagents (Takara), essentially as described^29^. RESISTΔC constructs were also generated by PCR to remove residues after D161 followed by Infusion cloning into the pLIP backbone. To generate retrovirus, GP2s at > 70% confluency in a 6-well plate were transfected with 0.5 μg of VSV-G and 3.5 μg of retroviral vector per well with Lipofectamine 2000. Retroviral vectors in this study (SINV HA-SP140, SINV-SP140) were derived from the self-inactivating retrovirus pTGMP (Addgene plasmid #32716, from the lab of Scott Lowe). Mouse SP140 or HA-SP140 codon-optimized cDNA was cloned into pTGMP with Infusion (Takara), modified to include a minimal CMV promoter driving SP140 constructs, followed by a PGK promoter driving a puromycin resistance cassette. Eighteen-20 hours after transfection, media on transfected cells was changed to 1 mL BMM media. Bone marrow was harvested as described above, and plated in BMM media without dilution. Retronectin (Takara) treated 6-well plates were generated according to manufacturer instructions. The following day (∼30 hours after changing media on transfected cells), virus was harvested from transfected cells by filtration of supernatant through a 0.45 μm filter and added to 1 e6/well of bone marrow in 4 mL total of BMM media. Plates were spun at 650 *x g* for 1.5-2 hours at 37°C. Three days after bone marrow harvest, BMMs were fed with 1.33 mL BMM media. BMMs were puromycin-selected on day 4 after BMM harvest with 2.75-5 μg/mL puromycin.

Puromycin kill curves were determined for every stock to identify the lowest concentration needed for BMM selection, and a non-transduced well or BMMs transduced with a retroviral vector lacking a puromycin resistance cassette were used to verify complete killing of non-transduced cells by puromycin. After puromycin selection (2 days), media was exchanged and BMMs were allowed to recover for 2-6 days before seeding. BMMs transduced with lentiviral pLIP constructs were pre-treated overnight-24 hours with 2.5 μg/mL doxycycline, then restimulated with 100 μg/mL DMXAA and fresh doxycycline. Cells were harvested at timepoints indicated in the figure legends for either RNA isolation or co-immunoprecipitation.

### HA-SP140 CUT&RUN and analysis

*Sp140^−/−^* BMMs were transduced with retrovirus encoding HA-SP140 or SP140 as described above, and stimulated for 8 hours with 100 μg/mL DMXAA. Half a million cells were input into CUT&RUN^66^ in biological triplicates using the Epicypher CUTANA^TM^ ChIC/CUT&RUN kit (Epicypher, 14-1048, v3), using 0.5 μg of rabbit anti-HA monoclonal antibody (Cell Signaling Technologies, C29F4) with *E. coli* genomic DNA spike-in. Non-transduced B6 BMMs (0.5 e6) were also put into CUT&RUN with 0.5 μg of rabbit isotype control IgG (Epicypher, 13-0042), with a single biological replicate. CUT&RUN was carried out on isolated nuclei according to kit instructions. Libraries were prepared using the Epicypher CUTANA^TM^ CUT&RUN library prep kit (Epicypher, 14-1001) according to kit instructions, then sequenced on an Illumina Novaseq flow cell with 250 bp paired-end reads for ∼6 million reads per sample. Adapters and low-quality reads were trimmed using BBDuk v38.05 using options ‘ktrim=r k=23 mink=11 hdist=1 maq=10 tpe tbo qtrim=r trimq=10’. Trimmed reads were aligned to the mm10 assembly using BWA-MEM v0.7.15, and only uniquely mapping reads with a minimum MAPQ of 10 were retained. Fragments aligning to the mitochondrial genome were removed. Peak calling was performed using complete and size subsetted alignment files with MACS2 v2.1.1 with paired-end options ‘--format BAMPE –-pvalue 0.01 –-SPMR –B –-call-summits’. Bigwig files were prepared from the MACS2 normalized bedgraph files using bedGraphToBigWig v4. MACS2 peak scores, the normalized number of sequence reads that originate from a bound genomic location, were output for HA-SP140 peaks.

### Cistrome and GREAT analysis

Replicate MACS2 CUT&RUN peak files were merged, then controls (IgG and SP140) were subtracted from the HA-SP140 peak file using bedtools intersect (v2.28.0)^67^ to output a file of SP140 peaks. This SP140 peak file was used as input in the cistrome toolkit data browser (http://dbtoolkit.cistrome.org/)^68^ looking for significant binding overlap of histone marks and variants in mm10. The SP140 peak file was also used as input for Genomic Regions Enrichment of Annotations Tool (GREAT)^69^.

### Recombinant protein expression and purification

ANXA2-S100A was produced and purified in E. coli BL21 (DE3) Star cells (Thermo Fisher Scientific) in LB medium at 20 °C as a fusion protein carrying an N-terminal His6-SUMO tag. Cells were resuspended in lysis buffer (50 mM HEPES, 500 mM NaCl,

25 mM Imidazole, pH 7.5) and lysed using a Branson Ultrasonics Sonifier SFX550. The lysate was cleared by centrifugation at 40,000 × *g* for 1 hr at 4 °C. The cleared lysate was loaded onto a 5 ml HisTrap column (Cytiva). The bound protein was eluted over a linear gradient with elution buffer (50 mM HEPES, 200 mM NaCl, 500 mM Imidazole, pH 7.5). The final step was size exclusion chromatography on a Superdex 200 26/600 column in a buffer containing 10 mM HEPES, 200 mM NaCl, 2 mM DTT, pH 7.5.

For expression of full-length RESIST, full-length human RESIST (UniProt ID: Q3ZCQ2) was inserted between the BamHI and XbaI restriction sites of the pLIB plasmid^70^ with a TEV (tobacco etch virus) protease-cleavable, N-terminal His6-MBP (maltose-binding protein) tag, and a C-terminal StrepII tag. The DNA sequence encoding the RNA-binding zinc fingers of human TTP (TZF; UniProt ID: P26651 residues Ser-102 to Ser-169) was inserted between the NdeI and XhoI restriction sites of the pnYC plasmid^71^ with a TEV-cleavable, N-terminal MBP tag, and a C-terminal StrepII tag. DNA constructs for the expression of the NOT9 module were previously described^72^.

Subsequently, full-length human RESIST with an N-terminal, TEV-cleavable His6-MBP tag and a C-terminal StrepII tag was expressed in Sf21 insect cells utilizing the MultiBac baculovirus expression system^73,74^ as previously described^75^. Briefly, Sf21 cells were grown to a density of 2×106 cells/ml at 27°C in Sf900II medium (Thermo Fisher Scientific), infected with the V1 generation His6-MBP-RESIST-StrepII baculovirus, and harvested 48 h after they stopped dividing. Harvested cells were resuspended in ice-cold protein buffer (50 mM HEPES pH 7.5, 150 mM NaCl, 5% v/v glycerol, 20 mM CHAPS, 25 mM imidazole) and lysed by sonication. The lysate was clarified by centrifugation at 40,000 g for 40 min at 4 °C, filtered through a 0.45-micron nylon filter, and loaded on a 5 ml nickel-charged HisTrap column (Cytiva). Contaminants were removed by washing with lysis buffer supplemented with 40 mM imidazole, and His6-MBP-RESIST-StrepII was eluted in lysis buffer supplemented with 250 mM imidazole.

The eluted protein was further purified by size exclusion chromatography on a HiLoad Superdex 200 16/600 column (Cytiva) in a buffer containing 50 mM HEPES pH 7.5, 150 mM NaCl, 5% v/v glycerol, 20 mM CHAPS. Peak fractions were pooled,

concentrated with a centrifugal filter, flash-frozen in liquid nitrogen, and stored at –80 °C. The RNA-binding zinc fingers of TTP (TZF) were expressed in E. coli BL21(DE3) Star cells (Thermo Fisher Scientific) in autoinduction media^76^ at 20 °C overnight as a fusion protein carrying an N-terminal, TEV-cleavable MBP tag, and a C-terminal StrepII tag.

Harvested cells were resuspended in protein buffer (50 mM HEPES pH 7.5, 300 mM NaCl, 10% w/v sucrose) and lysed by sonication. The lysate was clarified by centrifugation at 40,000 g for 40 min and loaded on a 1 ml StrepTrap XT column (Cytiva). Contaminants were removed by washing with high salt buffer (50 mM HEPES pH 7.5, 1 M NaCl, 10% w/v sucrose) before elution with lysis buffer supplemented with 50 mM biotin. Eluted protein was further purified by size exclusion chromatography on a Superdex 200 26/600 column (Cytiva) in protein buffer supplemented with 2 mM DTT.

The peak fractions were then pooled, concentrated with a centrifugal filter, flash-frozen in liquid nitrogen, and stored at –80 °C. Finally, The NOT9 module was prepared as previously described^47^.

### StrepTactin pull-down assay

StrepII-tagged MBP, as well as StrepII-tagged and SUMO-tagged SMARCA3 (residues 26-39) were produced in E. coli BL21 (DE3) Star cells (Thermo Fisher Scientific) grown in auto-induction medium overnight at 37 °C. Cells were resuspended in lysis buffer

(50 mM HEPES, 500 mM NaCl, pH 7.5) and lysed using a Branson Ultrasonics Sonifier SFX550, the lysate was then cleared by centrifugation at 40,000 × *g* for 1 hr at 4 °C. Purified ANXA2R was incubated with StrepTactin Sepharose resin (Cytiva, 28935599). After a 1-hr incubation beads were washed twice with 50 mM HEPES, 500 mM NaCl, pH 7.5, 0.03% Tween, once with 50 mM HEPES, 500 mM NaCl, pH 7.5, and once with binding buffer (50 mM HEPES, 200 mM NaCl, pH 7.5). Purified ANXA2-S100A was added to the bead-bound proteins. After a 1-h incubation, beads were washed four times with binding buffer and proteins were eluted with 50 mM biotin in binding buffer. The eluted proteins were analyzed by SDS-polyacrylamide gel electrophoresis followed by Coomassie blue staining.

For pulldowns of full-length human RESIST with CCR4-NOT subunits, purified His6-MBP-RESIST-StrepII or His6-MBP-StrepII were immobilized as bait via the C-terminal StrepII tag on streptavidin agarose resin prepared in-house. Two hundred fifty pmol of bait protein was incubated for 1 h in pull-down buffer (50 mM HEPES pH 7.5, 200 mM NaCl, 0.03% v/v Tween-20) at 6 °C under constant agitation. Unbound protein was removed following two washes with pull-down buffer, and 500 pmol of NOT9 module was incubated for 1 h with the bead-bound protein. Finally, the beads were washed three times with a pull-down buffer, and the bound proteins were eluted using a pull-down buffer supplemented with 50 mM biotin. Eluted proteins were analyzed using SDS-PAGE, followed by Coomassie blue staining.

### BlaER1 transduction and stimulation

Lentivirus was generated from HEK293T cells transfected with pLIP constructs (either mCherry or human RESIST either untagged or with N/C-terminal HA tags) as described for BMM transduction above. Virus was overlaid onto BlaER1 cells, which were subsequently puromycin selected and differentiated as described^29^. After 5 days of differentiation, BlaER1 cells were stimulated with 2.5 μg/mL doxycycline and fresh cytokines overnight. Media with new cytokines, fresh doxycycline, and ADU-S100 at a final concentration of 5 μg/mL (Aduro) was added the next day. At indicated timepoints, adherent and non-adherent cells were harvested and either lysed in TRK lysis buffer (Omega Biotek Total RNA kit) for RNA isolation or prepared for co-immunoprecipitation (below). RNA was isolated as quickly as possible from lysates as described above, and converted to cDNA with Superscript reagents and RNAse-inhibitors as described above.

### Co-immunoprecipitation

For IP of RESIST from BMMs, BMMs were transduced with RESIST constructs or mCherry and treated with doxycycline, followed by DMXAA treatment, as described above. Cells (0.6 e6 – 2.4 e6) were harvested at timepoints indicated in figure legends, and lysed on ice in 300-600 μL lysis buffer (50 mM Tris-HCl pH 7.5, 0.2% NP-40, 5% glycerol, 100 mM NaCl, 1.5 mM MgCl_2_, 1X protease inhibitor cocktail) for 30 minutes. Lysate was clarified by centrifugation at 18213 *x g* for 30 minutes at 4°C and quantified by BCA to ensure approximately equal amounts of input protein. One tenth of clarified lysate was diluted in Laemmli buffer for input sample. Supernatant was incubated with anti-HA magnetic beads (ThermoFisher, 88836) rotating 3 hrs – ON at 4°C. Beads with lysate were then washed three times with lysis buffer (1 mL per wash), then immunoprecipitated proteins were eluted in 30-50 mL Laemmli buffer by boiling for 5 minutes. Samples were then analyzed by immunoblot.

For IP of FLAG-TTP from BMMs^77^, N-terminally FLAG-tagged mouse TTP constructs was cloned into pLIP as described above. Mouse FLAG-TTP was transfected into semi-confluent 10 cm plates of HEK293Ts with N-terminally HA-tagged mouse RESIST or an equivalent amount of mCherry using Lipofectamine 2000 according to manufacturer instructions. Six hundred ng of TTP construct was co-transfected with 6 μg mCherry, and 660 ng of TTP construct was co-transfected with 6 μg RESIST. After transfection, cells were treated with doxycycline to induce expression for 24 hours, then harvested in PBS. Cells were lysed in hypotonic lysis buffer consisting of 10 mM Tris-HCl pH 7.5, 10 mM NaCl, 2 mM EDTA, 0.5% Triton X-100, and protease inhibitors for 10 minutes on ice with vortexing. NaCl was then adjusted to 150 mM and samples were treated with 30 μg of RNase A on ice for 20 minutes with vortexing. Samples were clarified via centrifugation at 18000 *x* g for 30 minutes at 4°C, and 10% of sample was taken as input sample and diluted in Laemmli buffer. Sample was incubated with 50 μL of Sigma Anti-FLAG M2 agarose beads (Sigma Aldrich, M8823) equilibrated in lysis buffer with adjusted NaCl and resuspended in a total volume of 200 μL lysis buffer with adjusted NaCl per reaction. Samples were incubated with beads for 2 hours rotating at 4°C, then washed 5 times with 1 mL wash buffer (50 mM Tris HCl pH 7.5, 300 mM NaCl, 0.05% Triton X-100, protease inhibitors). Remaining liquid was completely aspirated from beads with a small-bore needle, and beads were resuspended in 2X Laemmli buffer diluted in wash buffer before analysis by immunoblot.

### Immunoblot

Samples diluted in Laemmli buffer were run on 4-12% Bis-Tris protein gels (Invitrogen) and then transferred to PVDF membranes at 35 volts for 90 minutes. Membranes were blocked in either Odyssey Licor PBS blocking buffer or in 2-5% non-fat dry milk diluted in TBST. Membranes were then probed with antibodies at 4°C overnight diluted in 5% BSA TBST. Antibodies used in this study were rat anti-HA (Roche, clone 3F10, 1186742300, 1:1000), mouse anti-actin (Santa Cruz Biotechnology, sc-4478, 1:1000), rabbit anti-CNOT1 (Cell Signaling Technologies, 44613S, 1:1000), rabbit anti-CNOT9 (Proteintech, 22503-1-AP, 1:500), rabbit anti-TTP (Millipore Sigma, ABE285, 1:1000), rabbit anti-CNOT11 (Sigma Aldrich, HPA069823, 0.4 μg/mL), rabbit anti-ZFP36L1 (Cell Signaling Technologies, 30894S, 1:1000), rabbit anti-ZFP36L2 (Abcam, ab70775, 1:1000), and rabbit-SP140 (Covance, as previously described^5^; 1:1000), and rabbit anti-FLAG for FLAG-TTP IPs (Thermo Fisher Scientific, PA1-984B, 1:1000).

### In vivo Legionella pneumophila infections

*Legionella pneumophila* infections were performed as described previously^5^. Briefly, JR32 Δ*flaA Legionella pneumophila* (from the lab of Dario Zamboni) was streaked from a frozen glycerol stock onto BCYE plates. A single colony was used to streak a ∼ 4 cm^2^ patch that was subsequently grown for 2-3 days. Bacteria were diluted in water and OD measured at 600 nm to determine bacterial concentration. Bacteria were then diluted to a final concentration of 2.5E6 bacteria/mL in sterile PBS. Mice were anesthetized with a mixture of xylazine and ketamine via intraperitoneal injection, then 40 μL of diluted bacteria was administered intranasally (final infectious dose of 10^5^ bacteria per mouse). At 96 hours post-infection, mice were sacrificed and lungs were homogenized in 5 mL of autoclaved MilliQ water. Lung homogenate was diluted and plated on BCYE plates, and CFUs were enumerated after 4 days of growth.

### AlphaFold structure predictions

AlphaFold-Multimer v2.3.2^78^ was run on equipment hosted by the Cal Cryo EM facility comprised of an Nvidia GPU and >72 TB of storage space. Mouse RESIST and CCR4-NOT amino acid sequences were from NCBI (RESIST: XP_006517870.1; CNOT1 M-HEAT: XP_036009857.1, with a start codon followed by residues 815-1007; CNOT11: NP_082319.1; CNOT1 N-MIF4G:, XP_036009857.1, residues 1-695; CNOT10: NP_705813.2; CNOT9: NP_067358.1). Briefly, Alphafold was run in multimer mode on RESIST with CNOT1 M-HEAT, CNOT9, or CNOT1 N-MIF4G/CNOT10/CNOT11 with default settings and max template date specified as 2023-01-01, and *--db_preset=full_dbs.* Output models were visualized in ChimeraX (1.6.1), and aligned using the matchmaker command. PAE plots and structures colored by pLDDT were visualized with PAEViewer^79^.

### Gene disruption in BMMs with Cas9:gRNA electroporation

BMMs were harvested in PBS and electroporated with Cas9 2 NLS nuclease (Synthego) complexed with gRNAs (Synthego, sgRNA EZ kits), and Alt-R® Cas9 Electroporation Enhancer (IDT, 1075916), in Lonza P3 buffer (Lonza, V4XP-3032) with buffer supplement as described^80^. Electroporation was performed with a Lonza 4D-Nucleofector Core Unit (AAF-1002B) using the program CM-137. Electroporated BMMs were immediately plated in BMM media. A half-media exchange was performed every 2 days until day 10 after bone marrow harvest, when BMMs were seeded for downstream assays. Human BlaER1 cells transduced with RESIST constructs or mCherry were similarly electroporated, and cultured in complete RPMI. Knockout efficiency was evaluated by immunoblot or by PCR of genomic DNA for targeted regions followed by ICE analysis (Synthego) as indicated in figure legends. *Gm21188* (*Resist1*) was genotyped with Primestar PCR reagents and primers F1: ATTGAGAAATCCGTTTGTAATGGG and R1: TAGGCGAATTTCGTGGCACA according to manufacturer instructions with an annealing temperature of 55°C, or using Q5 PCR reagents and primers F2: TTGAGAAATCCGTTTGTAATGGG, R2: GCCTTTCTCCGGATTCACGA, with cycling conditions: 98°C 3 min, 35 cycles of 98°C 10 s, 63.5°C 20 s, 72°C 1:05 min. *Gm36079* (*Resist2*) was genotyped with F: TGGTATTCTCTAGAGATAACATCACAGCACCTACTTACTCC, R: CCTCCCCTCGCCATCACTGCCTG, using Phusion GC PCR reagents and cycling conditions of: 98°C 30 s, then 30-35 cycles of 98°C for 10 s, 72°C for 15 s, 72°C for 1 min. Disruption of *Rc3h1* was determined by PCR (F: CACACTATGTGCTGACTGTATCTACAGAAG, R: TCCCCTCAGGTAAAACAGTGC, cycling: 98°C 30 s, then 30 cycles of 98°C 10 s, 60°C 5 s, 72°C 1 min) with Phusion GC PCR reagents. Disruption of *Rc3h2* was determined by PCR (Q5 PCR reagents with F2: AGGGCATAAGATGTTGCACAGA, R2: ACTGCTAACCCGAGCATCAG: and cycling of 98°C 3 min, then 35 cycles of 98°C 10 s, 60°C 20 s, 72°C 40 s). PCRs were cleaned by gel extraction, Ampure XP beads (Beckman Coulter), or treatment with FAST-AP and ExoI before submission for Sanger sequencing. Guide RNA sequences used in this study were:

*Gm21188/Gm36079* gRNA 1: GCUGGGCCUCUUGCACCAGA

*Gm21188/Gm36079* gRNA 2: CGACGAUGGCGGUGACUACC

*Cnot1* gRNA 1: UGUGAAUCGGCACGGUCCUG

*Cnot1* gRNA 2: ACUCAUUCAGGAUUAACAGA

*Cnot11* gRNA 1: UCCAUCAAGGCAAUCUGGCG

*Cnot11* gRNA 2: GCUGAGCAUCAUCUCGGAGG

*Cnot9* gRNA 1: CAUUGCAAACUCUGUUAGAC

*Cnot9* gRNA 2: GCCUACUGCACUAGCCCAAG

*Zfp36* gRNA 1: CAUGACCUGUCAUCCGACCA

*Zfp36* gRNA 2: CUUCAUCCACAACCCCACCG

*Zfp36l1* gRNA 1: AAAAAUGGUGGCGGACACGA

*Zfp36l1* gRNA 2: ACGGGCAAAAGCCGAUGGTG

*Zfp36l2* gRNA 1: CAAGAAGUCGAUAUCGUAGA

*Zfp36l2* gRNA 2: GAGAGCGGCACGUGCAAGUA

*Rc3h1* gRNA: CAAAUGGGCAAGCCUUACGG

*Rc3h2* gRNA: UCGGUGAAGUUUAUUCAAGC

### TTP Electrophoretic mobility shift assays (EMSA)

The substrate RNAs were generated via in vitro transcription (IVT). For the IFNB1 WT (wild-type) RNA substrate, the AU-rich element (ARE) in the 3′UTR of the IFNB1 mRNA (GenBank ID NM_002176.4; nucleotides 740 to 825) was synthesized as a gene fragment (Azenta) with an upstream T7 promoter and 17 random nucleotides downstream. All adenosine residues between nucleotides 758 and 825 were mutated to cytosine for the IFNB1 MUT (mutated) RNA substrate. The gene fragments were amplified by PCR, and the purified PCR products were utilized as templates for IVT using the HiScribe T7 High Yield RNA Synthesis Kit (NEB). IVT products were separated via size exclusion chromatography on a Superdex 200 increase 10/300 gl in buffer containing 10 mM HEPES pH 7.5, 200 mM NaCl. Fractions containing the intact RNA substrates were pooled, ethanol precipitated, and resuspended in RNase-free water.

EMSA (electrophoretic mobility shift assay) binding reactions contained 50 nM substrate RNA and 50 to 800 nM TZF protein. Reactions were carried out for 15 min at 37 °C in a buffer containing 20 mM PIPES pH 6.8, 10 mM KCl, 40 mM NaCl, 2 mM Mg(OAc)2, 3% v/v Ficoll 400, and 0.05% v/v NP-40. The RNA-protein complexes were analyzed by electrophoresis on a nondenaturing polyacrylamide gel in 0.5x TBE buffer, pH 8.3, at 10 V cm-1. Gels were stained in 0.5x TBE pH 7.5 with 1x SYBR Gold (Thermo Fisher) for 5 minutes before analysis. Images were quantified using FiJi^81^.

### Viral infections of BMMs

Day 7 BMMs were generated as described above, and 250,000 cells were infected viruses at indicated multiplicities of infection (MOIs) in a non-TC treated 12 well plate. For MCMV-GFP and MHV68-GFP, cells were infected for 3-4 hours in serum-free RPMI supplemented with pen/strep and glutamine in a low volume of inoculum. MCMV-GFP and MHV68-GFP were a gift of Laurent Coscoy, UC Berkeley, and Britt A. Glaunsinger, UC Berkeley. MHV68-GFP and MCMV-GFP titer was estimated by infection of 3T3 cells, and calculated by the assumption that a viral dilution resulting in 100% of infected 3T3 cells corresponds to an approximate MOI of 5. For Sendai-GFP infections (ViraTree), cells were infected for 1.5 hours in serum-free RPMI supplemented with pen/strep and glutamine in a low inoculum volume. After infection, media was replaced and BMMs were cultured for an additional 20-24 hours before cells were harvested in PBS, stained with Ghost Dye Far Red 780, fixed with the BD Cytofix Cytoperm kit according to kit instructions, then washed and analyzed by flow cytometry. Data were analyzed with FlowJo.

### Immunofluorescence (IF) microscopy

BMMs were seeded on glass coverslips between 0.5 e6 – 1 e6/slip in BMM media lacking antibiotics and rested overnight. BMMs were stimulated with 100 μg/mL DMXAA for 8 hours, fixed in 4% freshly prepared paraformaldehyde (Electron Microscopy Sciences) for 10 minutes at room temperature, then permeabilized in freshly made 0.2% Triton X-100 and 0.2% BSA in PBS on ice for 10 minutes. Coverslips were washed 3 times with PBS and then blocked in goat serum and FC-block (TruStain FcX™ PLUS, anti-mouse CD16/32) for one hour, then incubated overnight at 4°C in primary antibody diluted in PBS with 1% Tween-20 and 1% BSA. Primary antibodies and dilutions were mouse anti-PML Millipore Sigma, 05-718, 1:100; rat anti-HA, Roche, 11867423001, 1:200; rabbit anti-Fibrillarin, Abcam, ab166630 1:100. Coverslips were washed 3 times in PBS, then incubated in secondary antibody diluted 1:1000 in PBS with 1% Tween-20 and 1% BSA for 2-3 hours at room temperature. Secondary antibodies used were donkey anti-rat Alexa Fluor 488, Invitrogen, A21208; goat anti-mouse Alexa Fluor 647, Invitrogen, A21236; goat anti rabbit 647, Life technologies, A21244. After 3 PBS washes of coverslips, nuclei were stained with DAPI at 1 μg/mL in PBS for 10 minutes at room temperature. Coverslips were mounted in Vectashield mounting medium (Vector Laboratories, H-1000-10). Coverslip edges were then sealed with clear nail polish before imaging on a Zeiss LSM 880 NLO AxioExaminer at 63x magnification.

### IF image processing and quantification

For independent experiments as indicated in figure legends, Z stacks were processed by Imaris File Converter v10.0.1 followed by Imaris Stitcher v9.9.1. Images were Gaussian filtered (0.132 μm) and screenshots were generated for figures. Surfaces for DAPI, HA-SP140, PML, and fibrillarin were generated in Imaris using split-touching of 1 μm for HA-SP140, PML, and fibrillarin surfaces and 5 μm for DAPI surfaces. Surface statistics were then exported. Surfaces were first filtered as DAPI+ (within 2 standard deviations below average fluorescence intensity of DAPI surfaces), then filtered by size (over 0.2 μm^3^). HA-SP140 nuclear bodies were considered to overlap fibrillarin if fibrillarin intensity mean for an HA-SP140 surface was within 2 standard deviations of average fluorescence intensity for fibrillarin surfaces, and vice versa for fibrillarin surfaces overlapping with HA-SP140 surfaces. The same criteria for overlap were applied to HA-SP140 and PML surfaces. Mean fluorescence intensities of surfaces were calculated for over 100 nuclei for each independent experiment.

### Statistical analysis and schematics

All results except HA-SP140 CUT&RUN and AlphaFold predictions were repeated at least twice in independent experiments. Statistics and graphs for all experiments except RNA-seq, ATAC-seq, and CUT&RUN experiments were generated using GraphPad Prism (version 10.0.2). For data with two groups of comparison, *p* values (* = *p* < 0.05, ** = *p* < 0.005, *** = *p* < 0.0005, **** = *p* < 0.00005, ns = not significant) were calculated with two-tailed t-tests using Welch’s correction or two-way ANOVAs, as detailed in figure legends. For data with more than two comparison groups, ANOVA tests were used (* = *p* < 0.05, ** = *p* < 0.005, *** = *p* < 0.0005, **** = *p* < 0.00005, ns = not significant). We found that the residuals for our RT-qPCR data were not normally distributed, and therefore for this data we performed ANOVA tests on log10-transformed data, which generated more normally distributed residuals based on Q-Q plots and is therefore more appropriate for an ANOVA test. We used one-way Welch’s and Brown-Forsythe ANOVA tests without assuming data sphericity or equal variance, for data with multiple genotypes and one treatment condition, with a more conservative Dunnett’s T3 multiple comparison correction post hoc for log10-transformed data and less conservative FDR correction for multiple comparisons post-hoc (Q= 0.001, two-stage linear step-up procedure of Benjamini, Krieger, and Yekutieli) for all other data. Two-way ANOVA tests were performed for data with more than two comparison groups and/or multiple timepoints of measurement, with a full model including an interaction term, as we found that the effect of genotype varied across time. For two-way ANOVA tests, we did not assume sphericity, and used Tukey’s multiple comparison test post-hoc or Šidák’s multiple comparison correction as detailed in figure legends. For non-normally distributed data (*Legionella pneumophila in vivo* infections), a Mann-Whitney test was used, with * = *p* < 0.05, ** = *p* < 0.005, *** = *p* < 0.0005, **** = *p* < 0.00005, ns = not significant. The mean for all data was graphed, and replicates are individually represented by dots. Error bars indicate standard error. Data shown in each figure represent the provided source raw data; statistical test details are also provided in source data. Replicates in RT-qPCR, ELISA or viral infections represent separate wells within an experiment, while replicates in *Legionella pneumophila* infections represent individual mice from three combined experiments. Replicate numbers (*n)* is represented in the figures and source data. Briefly, each experimental group had between *n* = 2-6 replicates per group, as depicted in figures with individual dots, for *in vitro* experiments. For *in vivo Legionella pneumophila* infections, *n* = 14 mice for B6, *n* = 20 mice for *Sp140^−/−^*, and *n* = 13 mice for *Sp140^−/−^ Resist1^−/−^Resist2^−/−^*. Schematics were generated in Biorender.

## Data availability

HA-SP140 anti-HA CUT&RUN data is available at GEO accession: GSE269315. RNA-seq for *Ifnar^−/−^* and *Sp140^−/−^Ifnar^−/−^*BMMs is available at GEO accession: GSE269761. RNA-seq and ATAC-seq for B6 and *Sp140^−/−^*BMMs is available at GEO accession GSE269808 and GSE269811 respectively.

## Source code

Source code for all analysis scripts and pipelines is available at https://github.com/adziulko/The-SP140-RESIST-pathway.

## Author contributions

Conceptualization: K.C.W. and R.E.V. Data analysis: K.C.W., A.D., J.A., P.A., A.X.C., E.B.C. Investigation: K.C.W., A.D., J.A., F.P., A.X.C., G.Y.L., O.V.L., D.J.T., M.M.G., R.C., S.A.F., P.A., D.I.K., L.C., E.V., E.B.C., R.E.V. Writing: K.C.W., A.D., and R.E.V. Resources: A.L., H.D. A.Y.L., B.A.G., L.C., G.Y.L. Funding acquisition: K.C.W., R.E.V.

## Acknowledgements

We thank the Vance and Barton labs for input and advice, and thank Ella Brydon for technical assistance. We thank Dr. Jessie Li (UC Davis Bioinformatics) for preliminary analysis of ATAC-seq and CUT&RUN datasets, Kartoosh Heydari, Melaine Delcroix, and Harman Dhaliwal in the Berkeley CRL Flow Cytometry Facility, and Dr. Evan Witt for advice on statistical and bioinformatic analyses. R.E.V. is an Investigator of the Howard Hughes Medical Institute and a Bakar Faculty fellow, and research in his lab is supported by NIH grants AI075039, AI063302, and AI155634 and an Emerging Pathogens Initiative grant from HHMI. K.C.W. was supported by the National Science Foundation Graduate Research Fellowship Program. F.P., D.J.T, and E.V. were supported by the Intramural Research Program of the National Institutes of Health. F.P. was further supported by a Walter Benjamin postdoctoral fellowship from the German Research Foundation (Deutsche Forschungsgemeinschaft, Project number 531520533). S.A.F. was supported by a Postdoctoral Fellowship from EMBO (ALTF 617-2021) and the Swiss National Science Foundation (P500PB_206801).

## Extended Data

**Extended Data Figure 1.**
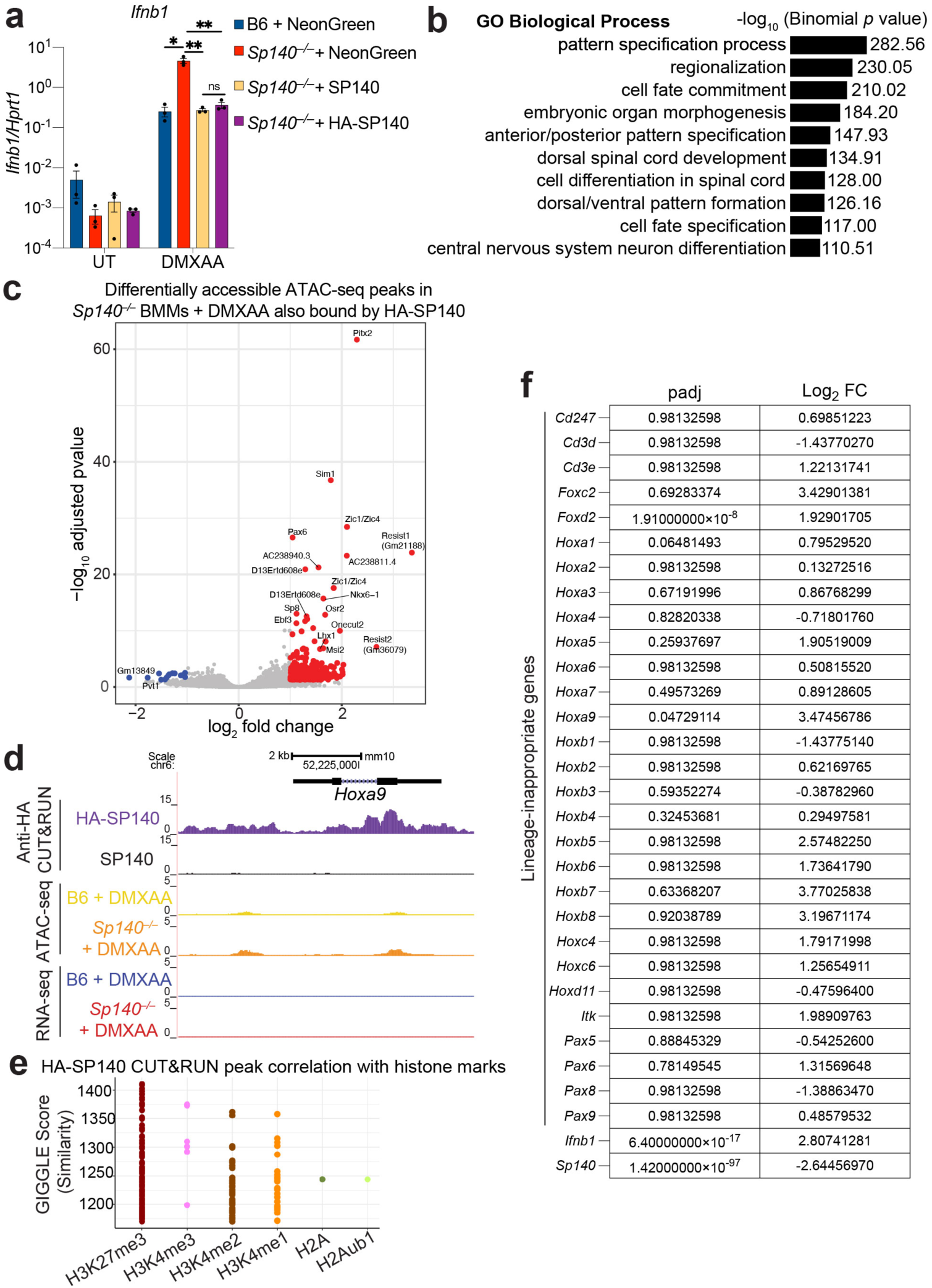
SP140 predominantly represses chromatin accessibility and binds genes involved in development, although these genes are not differentially expressed in the absence of SP140. a. RT-qPCR for *Sp140^−/−^* BMMs transduced with indicated retroviral constructs, treated with 100 μg/mL DMXAA for 8 hours. * = *p* < 0.05, ** = *p* < 0.005, *** = *p* < 0.0005, **** = *p* < 0.00005, ns = not significant, one-way ANOVA with Dunnett’s T3 post-hoc correction. b. Top 10 GO terms for genes bound by HA-SP140 in anti-HA CUT&RUN. c. Volcano plot of differentially accessible ATAC-seq peaks in DMXAA-treated *Sp140^−/−^* vs. B6 BMMs, filtered by genes that are also bound by HA-SP140 in anti-HA CUT&RUN. Blue dots indicate genes with log_2_ fold change < –1 and adjusted *p* (padj) < 0.05, and red dots indicate genes with log_2_ fold change > 1 and padj < 0.05. for differential ATAC-seq peak accessibility in *Sp140^−/−^* BMMs treated with DMXAA. d. Alignment of reads from HA-SP140 or untagged SP140 anti-HA CUT&RUN, and RNA-seq/ATAC-seq of *Sp140^−/−^* and B6 BMMs treated with DMXAA at *Hoxa9*. e. GIGGLE similarity score for overlap of HA-SP140 CUT&RUN peaks with publicly available ChIP-seq datasets for indicated histone marks^68^. f. Table of adjusted *p* (padj) and log_2_ fold change for “lineage-inappropriate SP140-regulated” genes^16^ from RNA-seq of DMXAA-treated *Sp140^−/−^* and B6 BMMs.

**Extended Data Figure 2.**
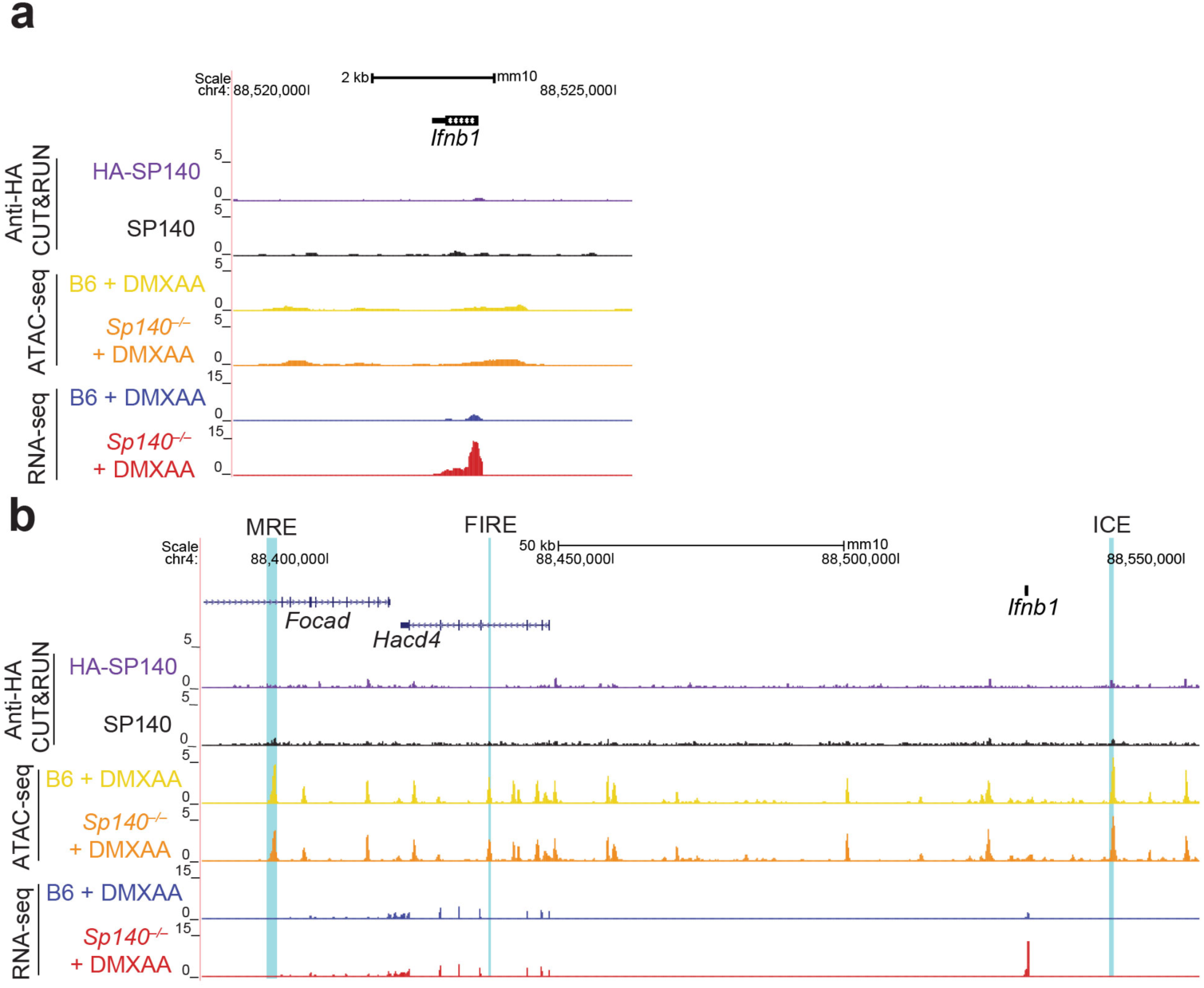
SP140 does not bind the *Ifnb1* locus or known regulatory elements. a. Alignment of reads at *Ifnb1* from HA-SP140 or untagged SP140 anti-HA CUT&RUN, ATAC-seq of DMXAA-treated *Sp140^−/−^* and B6 BMMs, and RNA-seq of DMXAA-treated *Sp140^−/−^* and B6 BMMs. b. Alignment of reads from HA-SP140 or untagged SP140 anti-HA CUT&RUN, and ATAC-seq/RNA-seq of *Sp140^−/−^* and B6 BMMs treated with DMXAA, at the *Ifnb1* regulatory elements ICE^31,32^, FIRE^30^, and the MRE^29^.

**Extended Data Figure 3.**
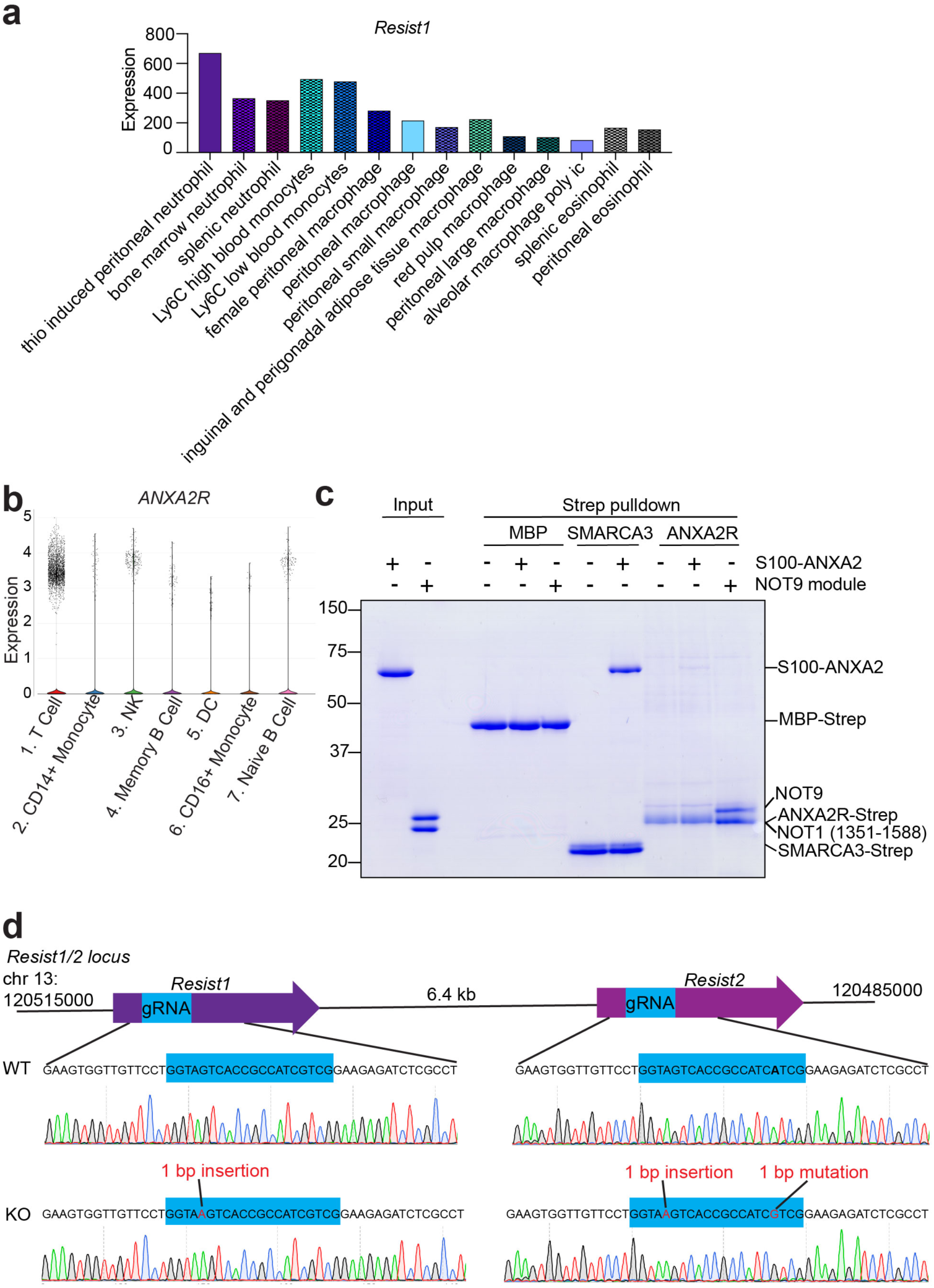
ANXA2R/RESIST expression in humans and mice, assessment of binding to Annexin 2, and generation of *Gm21188*/*Resist1^−/−^ Gm36079/Resist2^−/−^* mice. a. Expression values (DESeq2 normalized values) for *Gm21188* (*Resist1)* for all cell types with high expression (>80 for expression value). Data from immgen.org. b. Expression of *ANXA2R* in human PBMC single-cell RNAseq data from Immune Cell Atlas (data from https://singlecell.broadinstitute.org/single_cell/study/SCP345/ica-blood-mononuclear-cells-2-donors-2-sites?scpbr=immune-cell-atlas#study-summary) c. Pull-down assay of recombinant STREP-ANXA2R and STREP-SMARCA3 (residues 26-39) upon incubation with ANXA2-S100A. For gel source data, please see Supplementary Figure 1. d. Schematic of *Resist1/2 (Gm21188/Gm36079)* locus with protein coding sequences indicated with purple arrows and gRNA targeted sequence indicated in blue. *Resist2* contains a SNP within the gRNA targeting sequence (in bold). WT traces are indicated at guide-targeted regions for both *Resist1* and *Resist2*. Sequence traces from *Sp140^−/−^Resist1^−/−^Resist2^−/−^*mice (KO) are below with indicated mutations in *Resist1/2*.

**Extended Data Figure 4.**
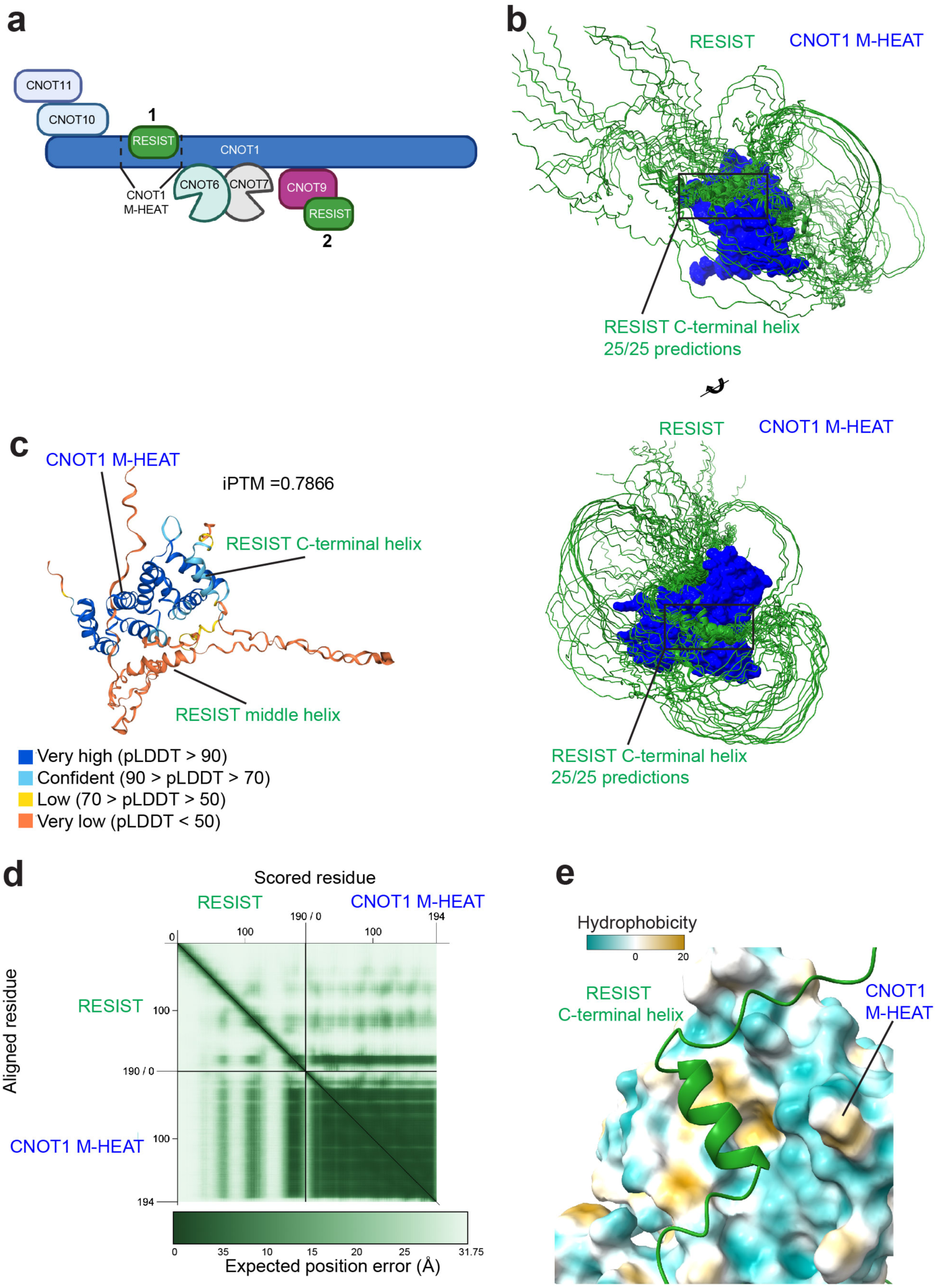
AlphaFold predictions suggest RESIST likely binds the CNOT1 M-HEAT domain. a. Schematic of CCR4-NOT subunit and RESIST complexes produced by Alphafold Multimer. b. Aligned AlphaFold models of RESIST with the CNOT1 M-HEAT domain. c. The top-scoring structural prediction of the RESIST and CNOT1 M-HEAT complex, colored by pLDDT. d. AlphaFold PAE plot of predicted RESIST and CNOT1 M-HEAT complex. Plot generated with PAEViewer^79^. e. Depiction of RESIST binding to a hydrophobic patch on the CNOT-1 M-HEAT domain (CNOT1 colored by hydrophobicity).

**Extended Data Figure 5.**
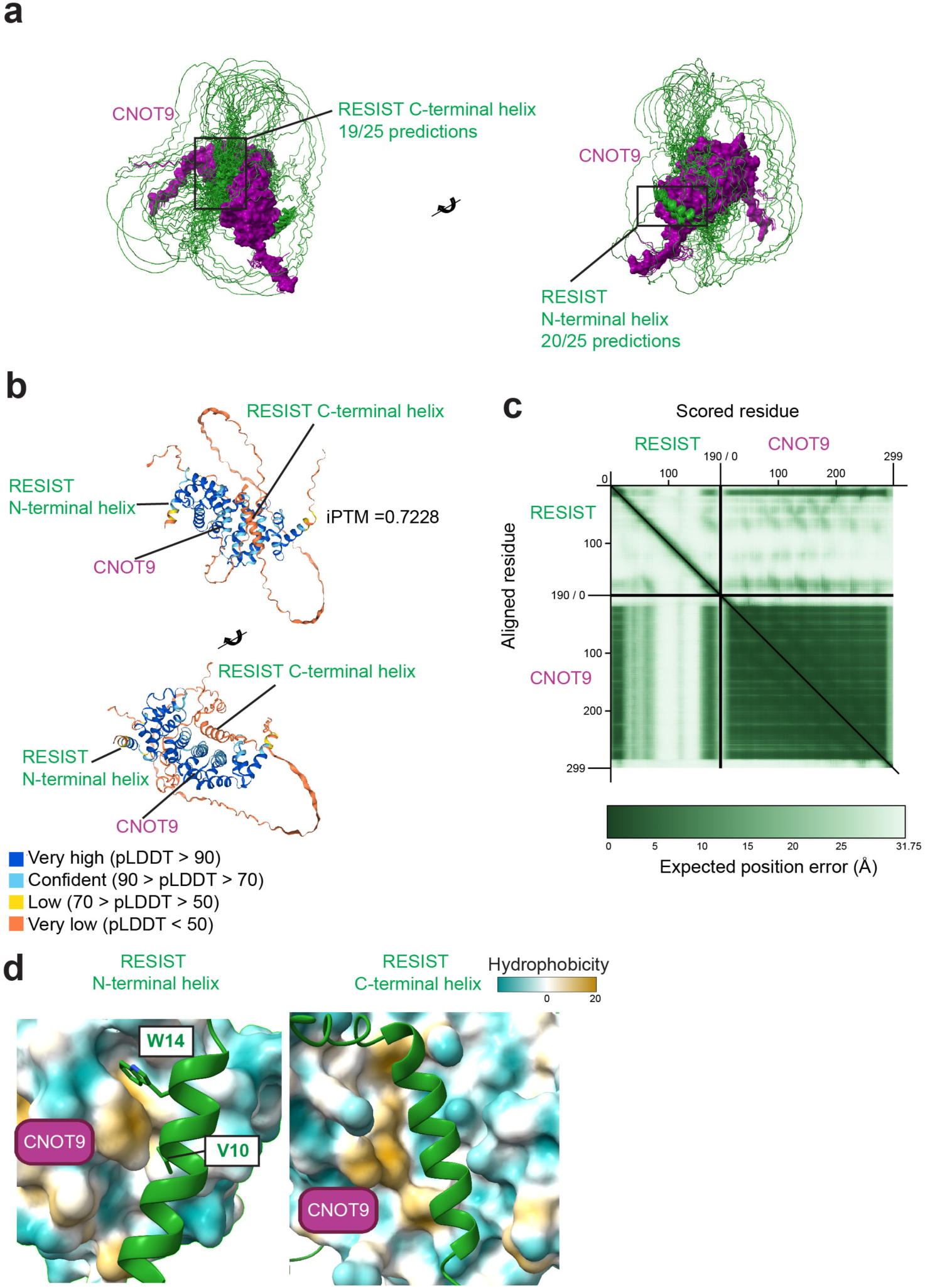
AlphaFold predictions suggest RESIST likely binds the CNOT9 subunit. a. Aligned structural predictions of RESIST interactions with CNOT9. b. Second-ranked Alphafold structural prediction of RESIST and CNOT9 complex colored by pLDDT. c. AlphaFold PAE plot of predicted RESIST and CNOT9 complex. Plot generated with PAEViewer^79^. d. Depiction of RESIST binding to multiple hydrophobic patches on CNOT9 (CNOT9 colored by hydrophobicity).

**Extended Data Figure 6.**
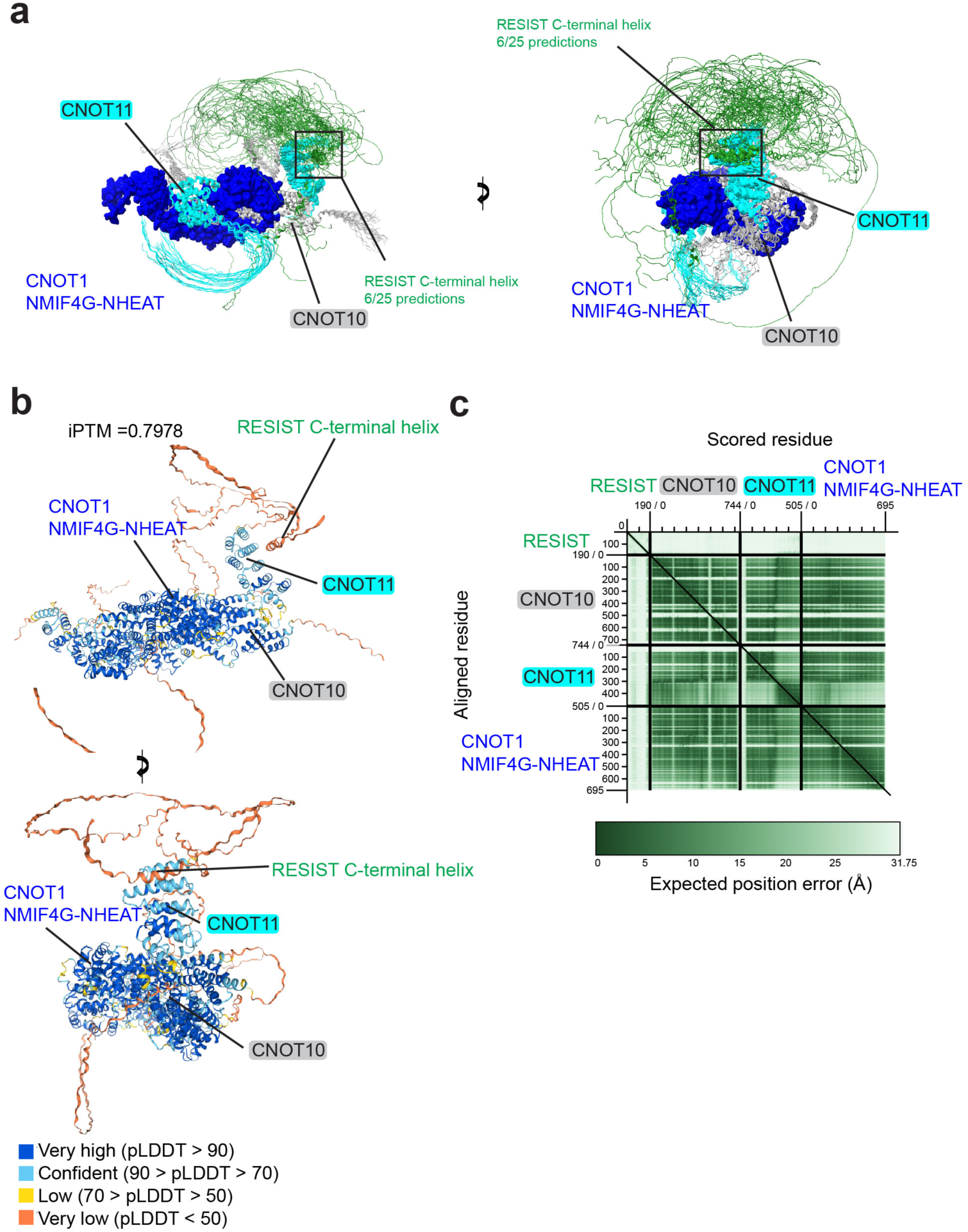
AlphaFold predictions suggest RESIST likely does not bind CNOT11. a. Aligned AlphaFold structure predictions of RESIST with the CNOT1 NMIF4G-NHEAT domains, CNOT10, and CNOT11. b. Highest-scoring AlphaFold structure prediction of RESIST and CNOT1/CNOT10/CNOT11 complex colored by pLDDT. c. AlphaFold PAE plot of predicted RESIST and CNOT1/CNOT10/CNOT11 complex. Plot generated with PAEViewer^79^.

**Extended Data Figure 7.**
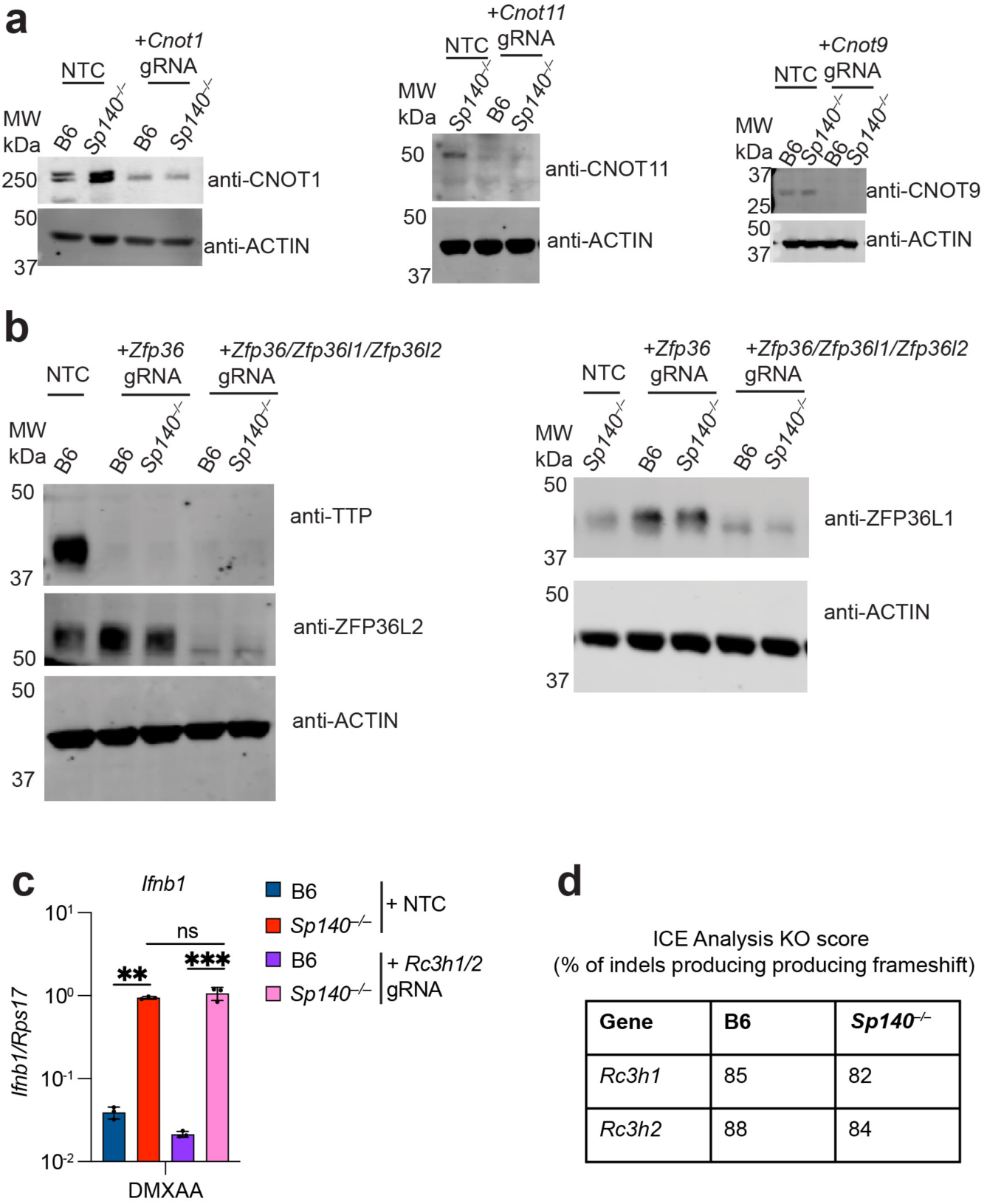
Assessment of CCR4-NOT subunit, TTP family, and ROQUIN1/2 knockout efficiency and role of ROQUIN1/2 in *Ifnb1* regulation in BMMs. a. Immunoblot of B6 or *Sp140^−/−^* BMMs electroporated with indicated gRNAs for indicated CCR4-NOT subunits for experiments shown in Fig. 4b. Actin blots represents loading controls. For gel source data, please see Supplementary Figure 1. b. Immunoblot of B6 or *Sp140^−/−^*BMMs electroporated with indicated gRNAs for indicated TTP family members for experiments shown in Fig. 4f after 8 hours of 100 μg/mL DMXAA. Actin blots represents loading controls. For gel source data, please see Supplementary Figure 1. c. RT-qPCR for *Ifnb1* in BMMs electroporated with NTC gRNAs or gRNAS targeting *Rc3h1*/*Rc3h2* (genes encoding ROQUIN1/2) after 8 hours of 100 μg/mL DMXAA treatment. * = *p* < 0.05, ** = *p* < 0.005, *** = *p* < 0.0005, **** = *p* < 0.00005, ns = not significant, one-way ANOVA with Dunnett’s T3 post-hoc correction. d. Knockout efficiency for *Rc3h1*/*Rc3h2* in BMMs electroporated with gRNAs targeting *Rc3h1* and *Rc3h2* for results shown in c.

**Extended Data Figure 8.**
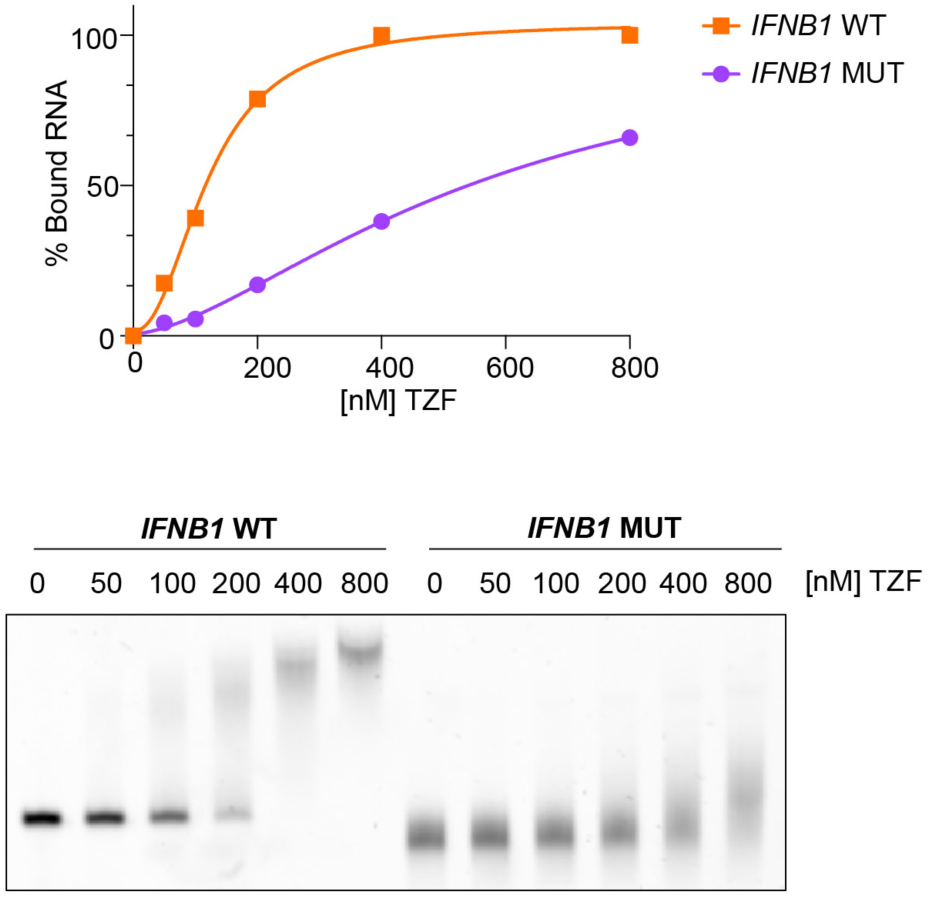
The TTP zinc finger domain (TZF) binds the *Ifnb1* 3’UTR in an ARE-dependent manner. Top: binding curve of TZF to SYBR-Gold labelled *Ifnb1* 3’UTR RNA for either WT (orange) or ARE mutant (purple), quantified from bottom. Bottom: representative electrophoretic mobility shift assay (EMSA) of TZF to ARE-WT or mutant *Ifnb1* 3’UTR. For gel source data, please see Supplementary Figure 1.

**Extended Data Figure 9.**
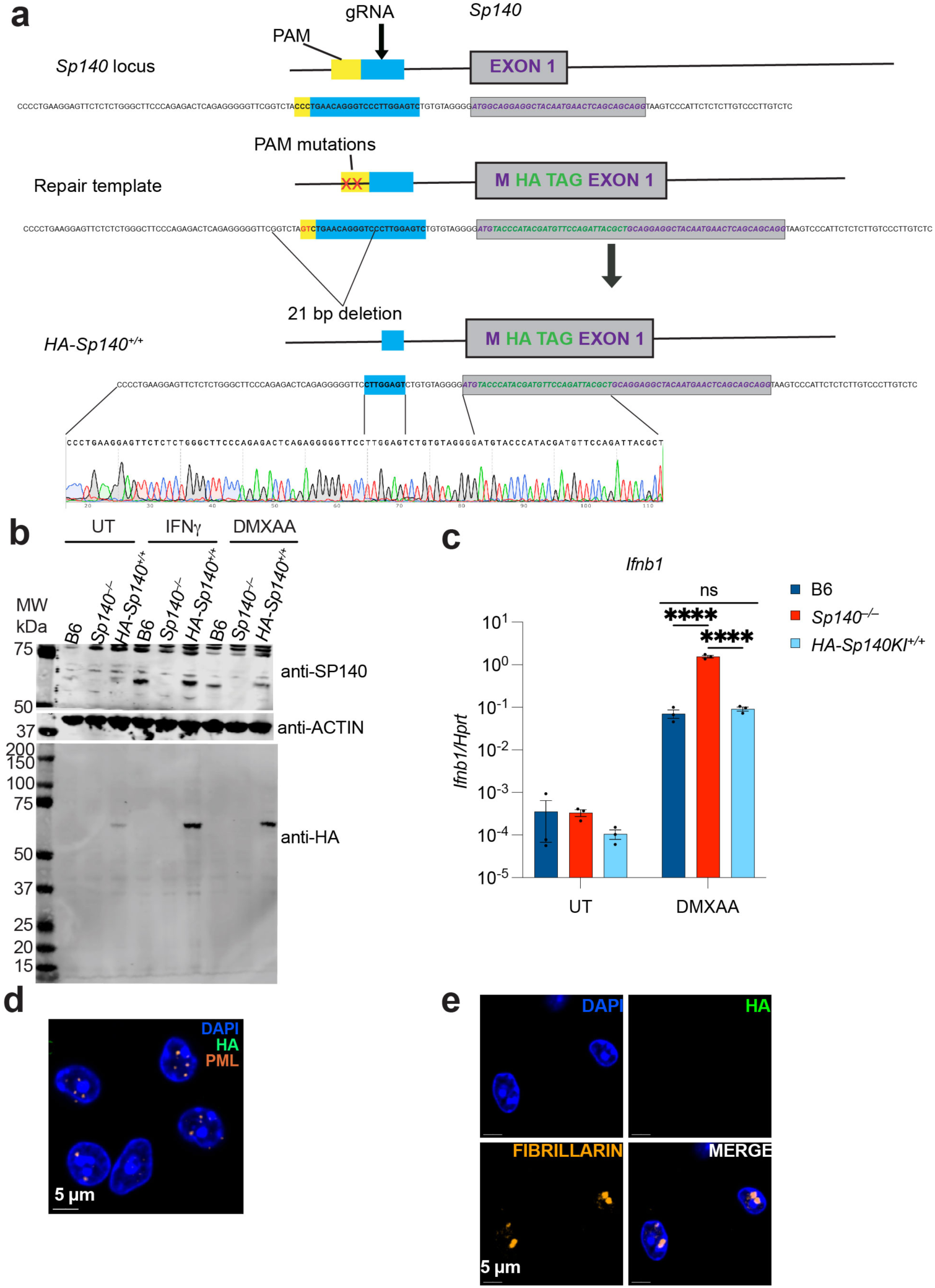
Generation and validation of *HA-Sp140* knock-in mice. a. Schematic of gene-targeting strategy to generate *HA-Sp140* knock-in mice and depiction of resulting *HA-Sp140^+/+^*founder line. b. Immunoblot of BMMs of indicated genotypes treated with 100 μg/mL DMXAA for 8 hours or 10 ng/mL IFNγ for 24 hours. Actin blot represents loading control for anti-SP140 blot, and sample processing control for anti-HA blot. For gel source data, please see Supplementary Figure 1. c. RT-qPCR for *Ifnb1* from BMMs of indicated genotypes after 8 hours of 100 μg/mL DMXAA treatment. * = *p* < 0.05, ** = *p* < 0.005, *** = *p* < 0.0005, **** = *p* < 0.00005, ns = not significant, one-way ANOVA with Dunnett’s T3 post-hoc correction. d. Immunofluorescence of B6 BMMs treated with 8 hours of 100 μg/mL DMXAA stained with anti-HA, anti-PML, and DAPI. Staining control for Fig. 5a. e. Immunofluorescence of B6 BMMs treated with 8 hours of 100 μg/mL DMXAA stained with anti-HA, anti-fibrillarin, and DAPI. Staining control for Fig. 5b.

**Extended Data Figure 10.**
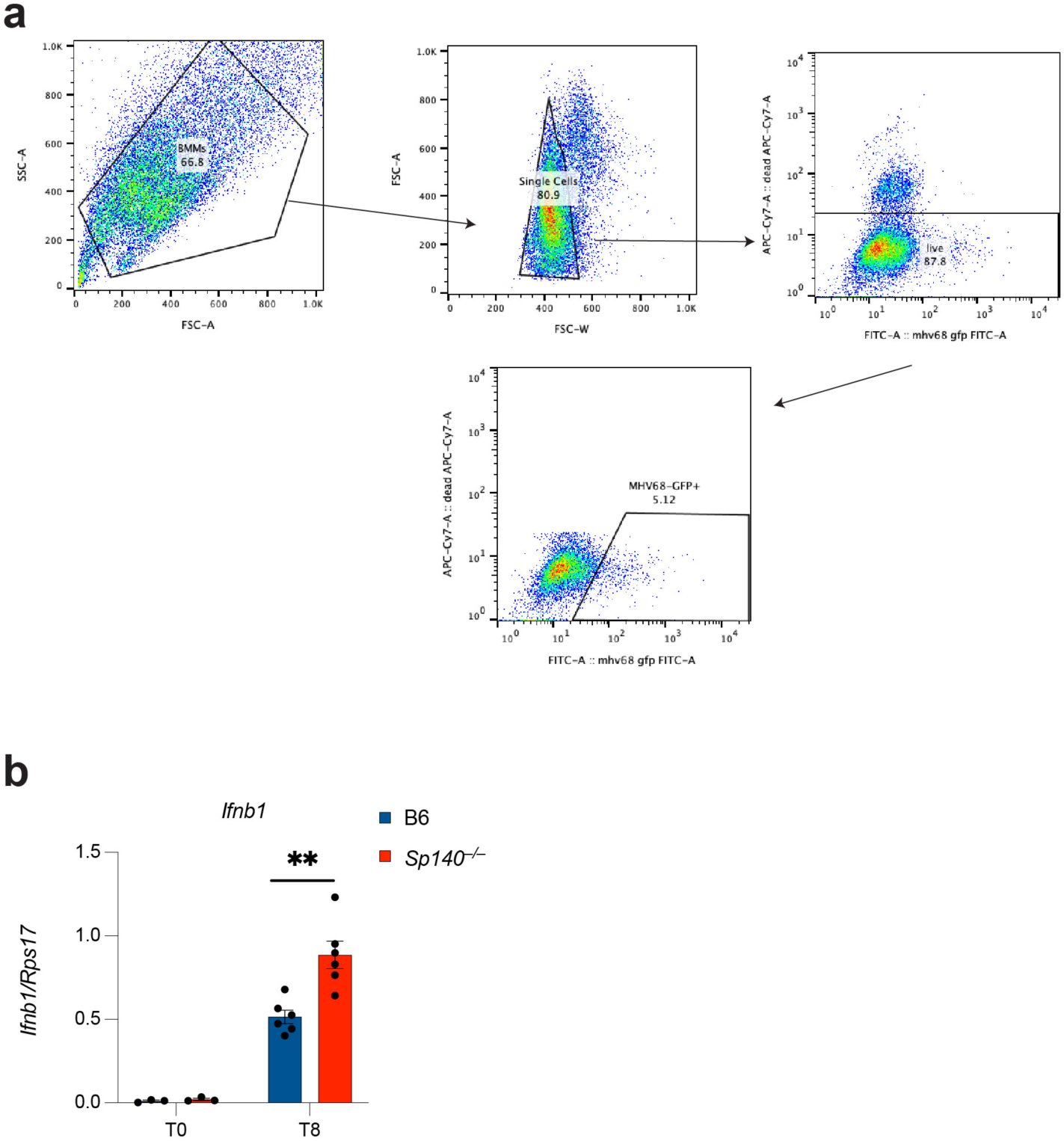
Gating strategy for BMMs infected with GFP-encoding viruses, and measurement of *Ifnb1* ranscripts in B6 and *Sp140^−/−^*BMMs upon infection with MHV68-GFP. a. Representative flow plots and gating strategy for BMMs infected with viruses encoding GFP. Experiment shown is for B6 BMMs infected with MHV68-GFP, MOI 3, for 24 hours. b. RT-qPCR of BMMs 8 hours after infection with MHV68-GFP, MOI of 1. * = *p* < 0.05, ** = *p* < 0.005, *** = *p* < 0.0005, **** = *p* < 0.00005, ns = not significant, two-tailed t-test with Welch’s correction.

