## Supplementary Figure 1 for "The SP140-RESIST pathway regulates interferon mRNA stability and antiviral immunity"

Figure 3a

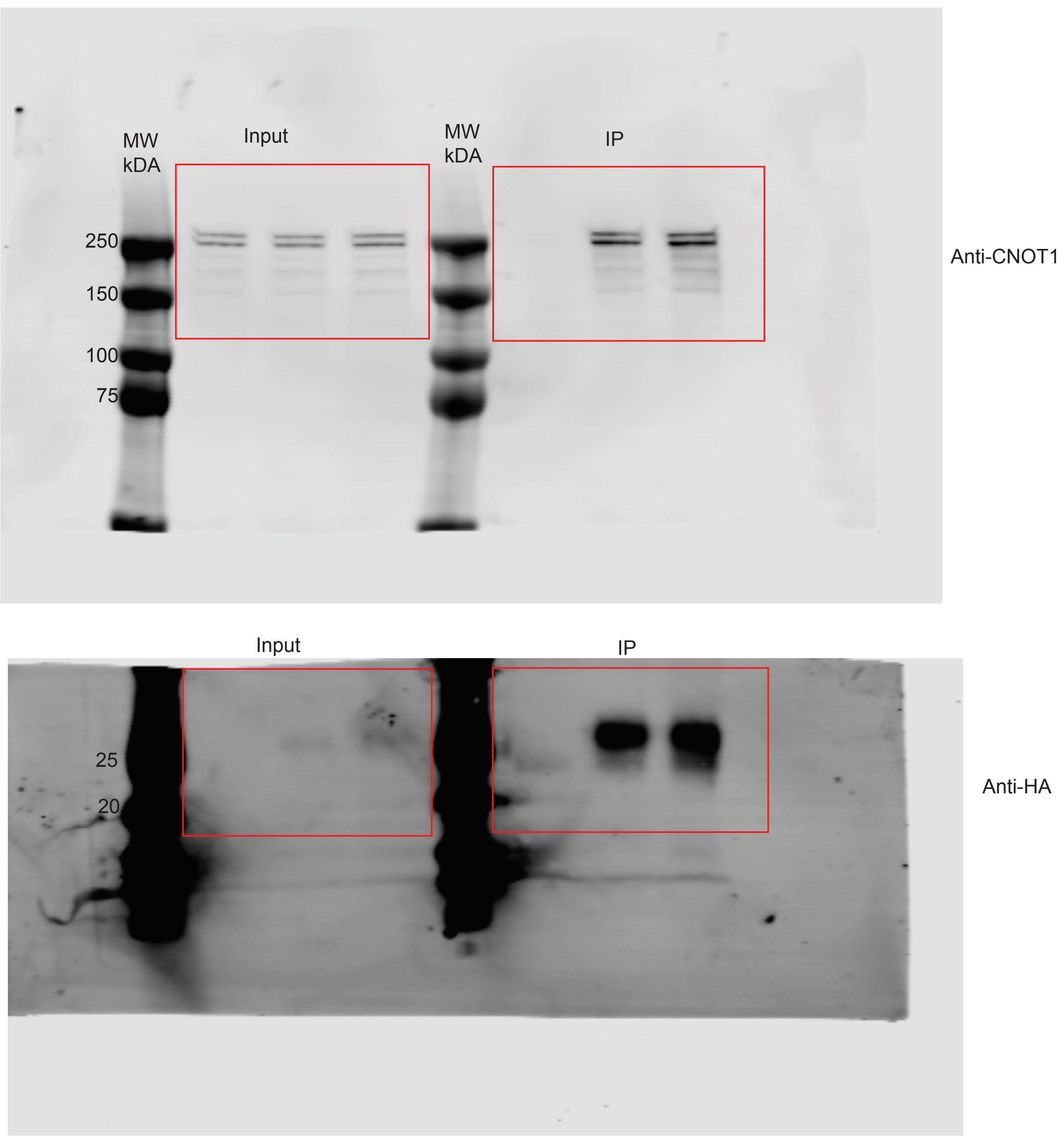
MW
kDA
Input
MW
kDA
IP
250
150
100
75
Anti-CNOT1
Input
IP
25
20
Anti-HA

Figure 4c

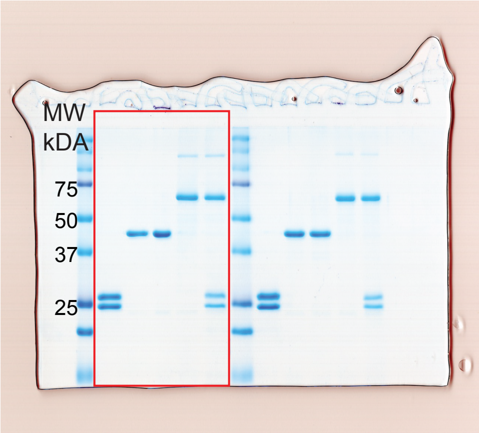


Figure 4e

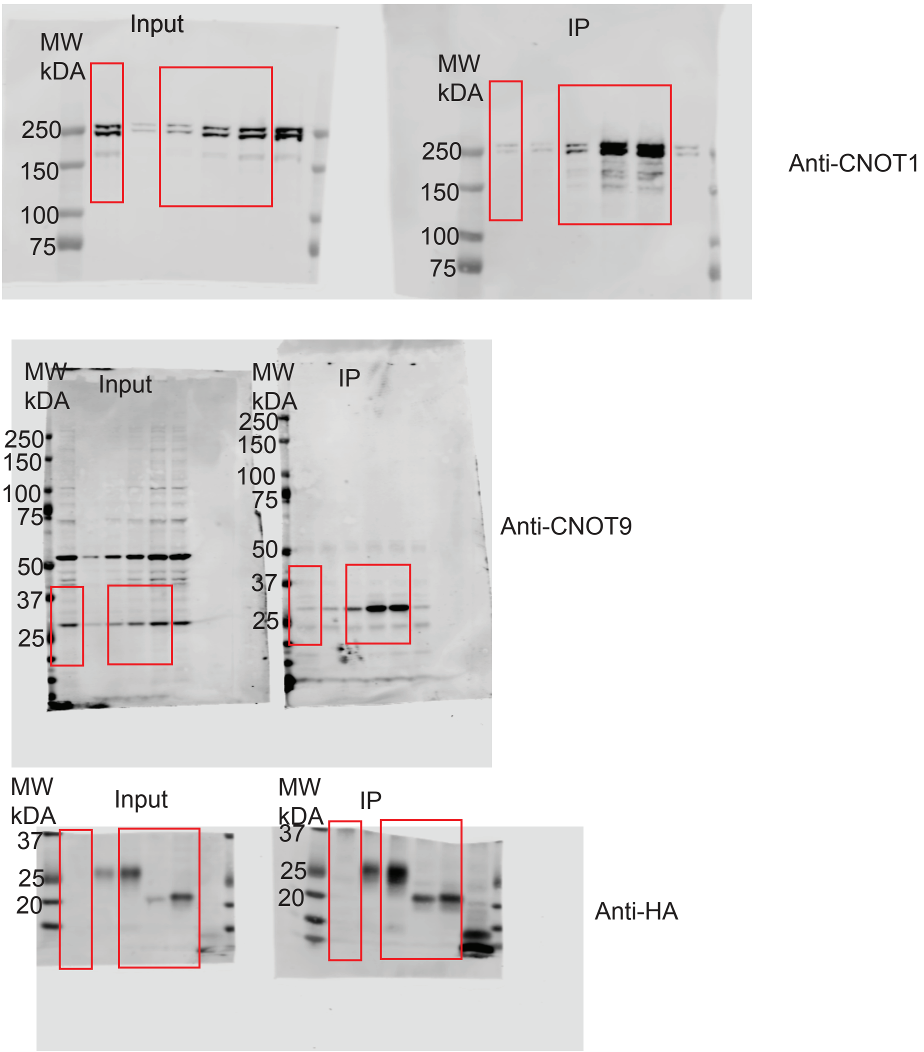

Figure 4g

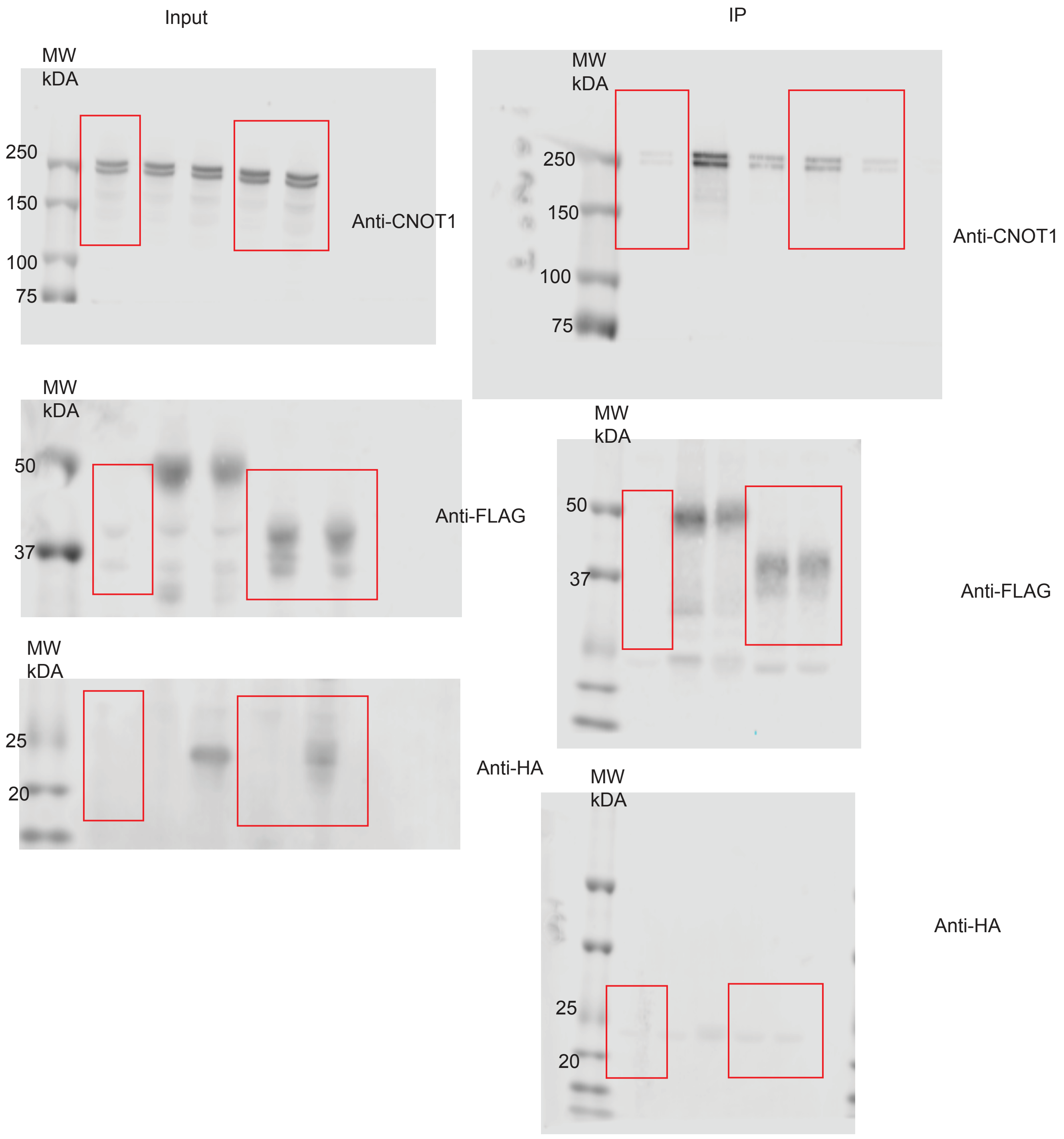
Input
IP
MW
kDA
250
150
100
75
Anti-CNOT1
MW
kDA
250
150
100
75
Anti-CNOT1
MW
kDA
50
37
Anti-FLAG
MW
kDA
50
37
Anti-FLAG
MW
kDA
25
20
Anti-HA
MW
kDA
25
20
Anti-HA

Extended Data Figure 3c

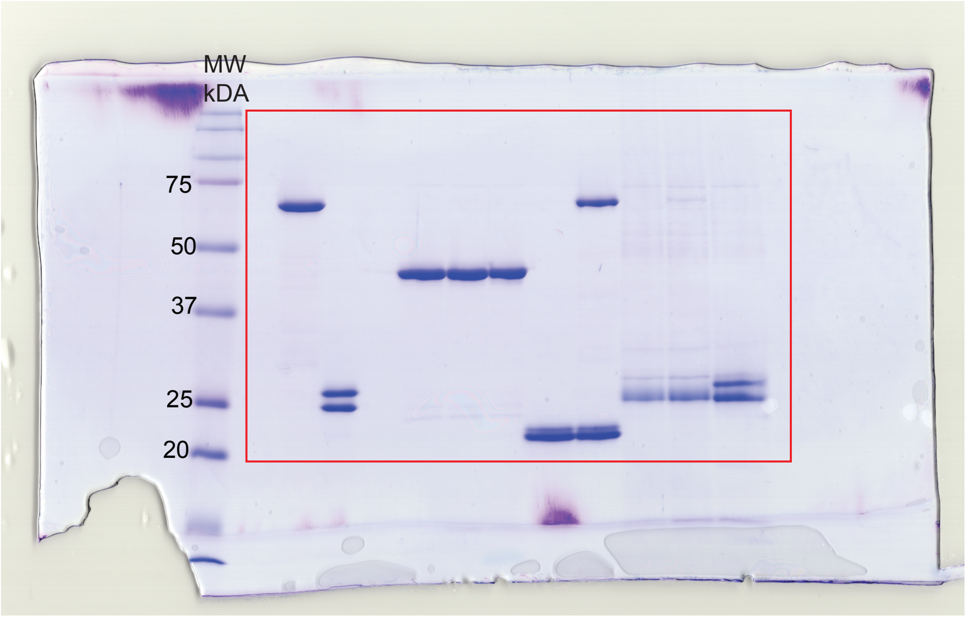

Extended Data Figure 7b

Left

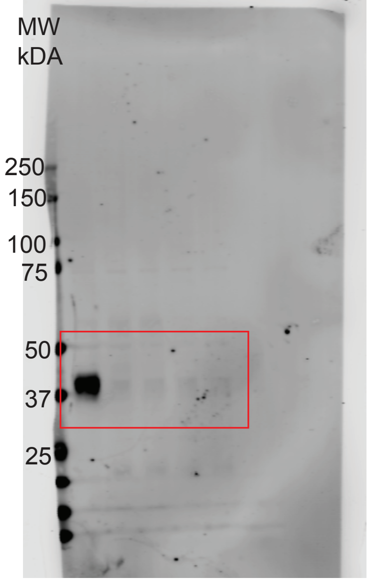


Anti-TTP

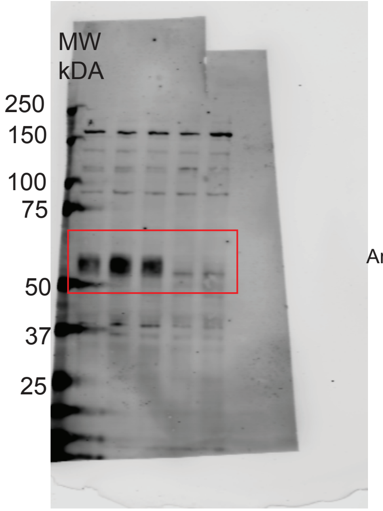


Anti-ZFP36L2

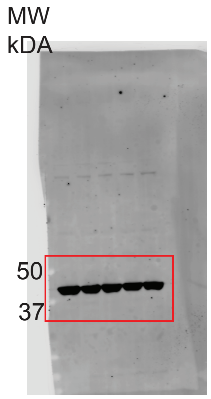


Anti-ACTIN for anti-ZFP36L2 and anti-TTP (loading control)

Right

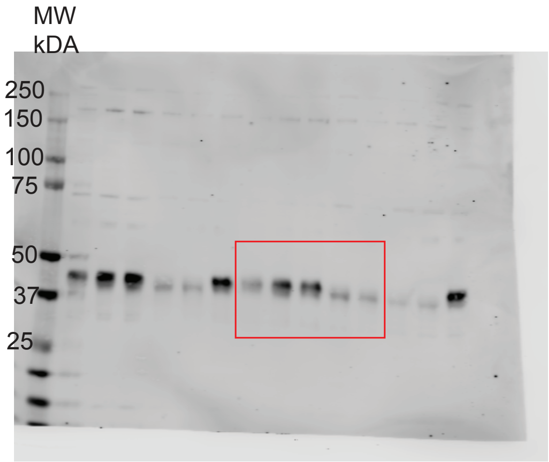


Anti-ZFP36L1

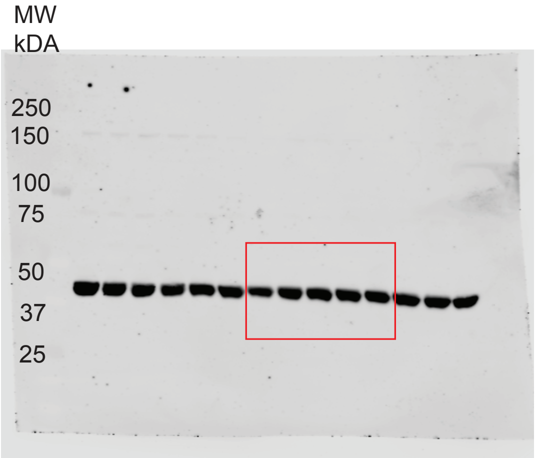


Anti-ACTIN for anti
-ZFP36L1
(loading control)

Extended Data Figure 7a

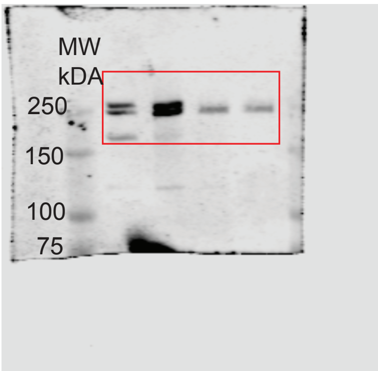


Anti-CNOT1

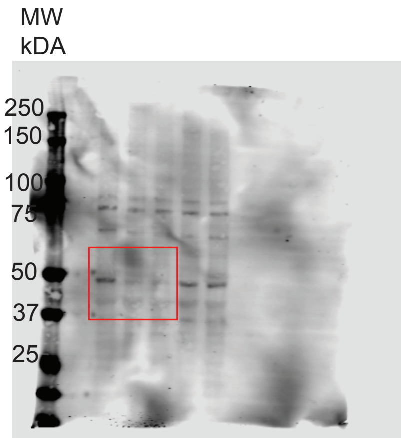


Anti-CNOT11

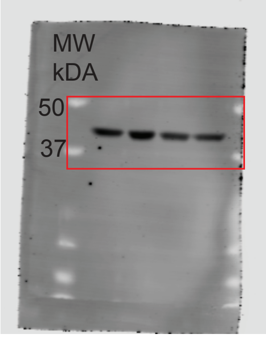


Anti-ACTIN for CNOT1 (loading control)

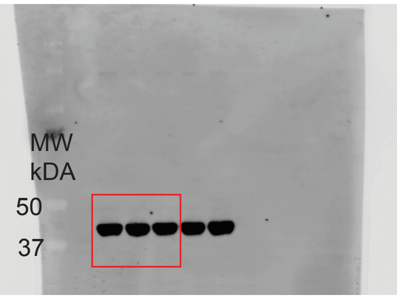


Anti-ACTIN for CNOT11 (loading control)

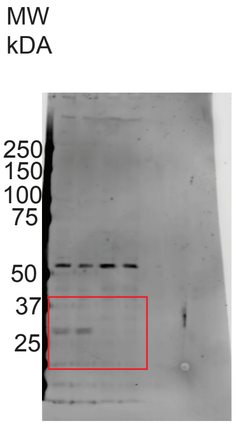


Anti-CNOT9

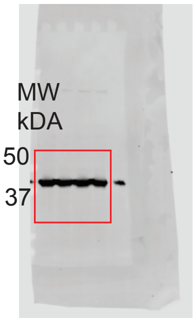


Anti-ACTIN for CNOT9 (loading control)

Extended Data Figure 8

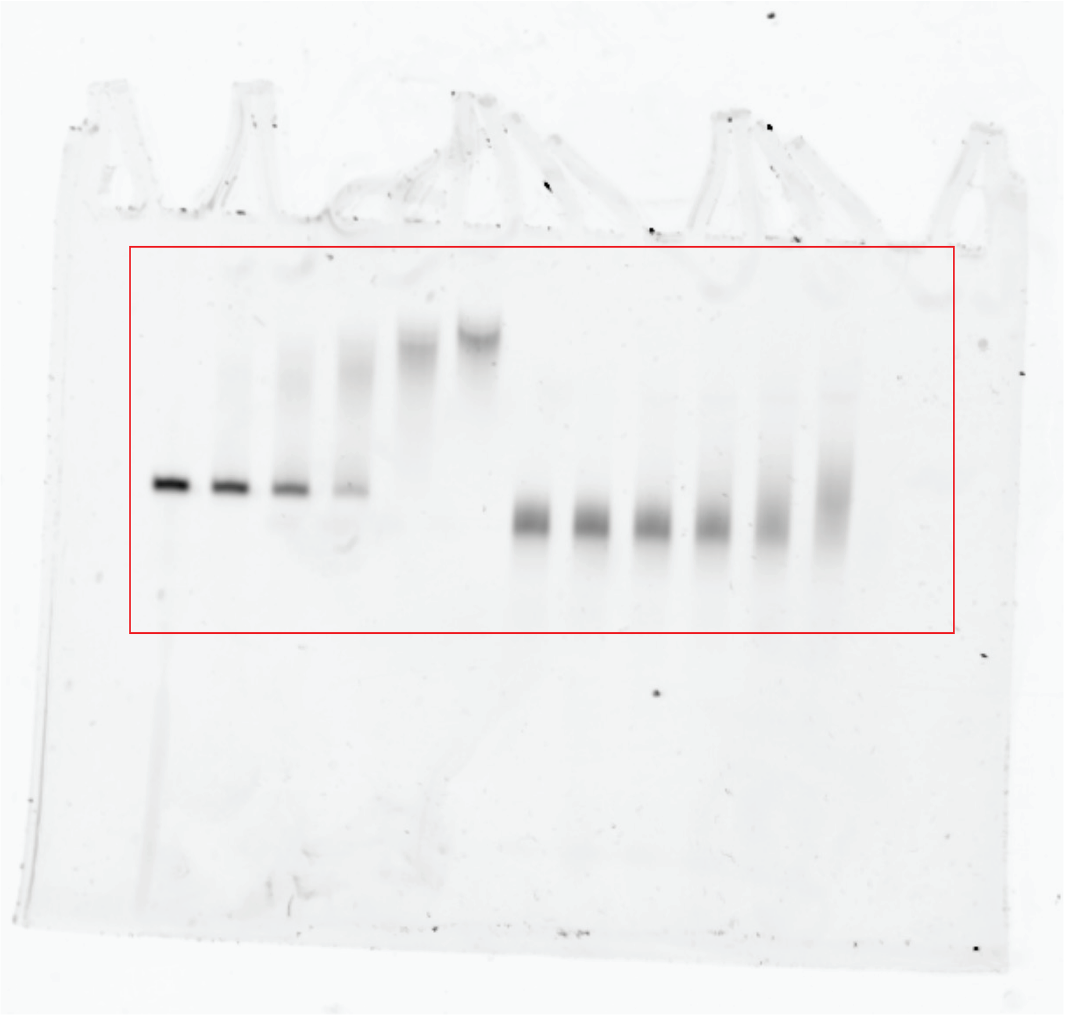

Extended Data Figure 9

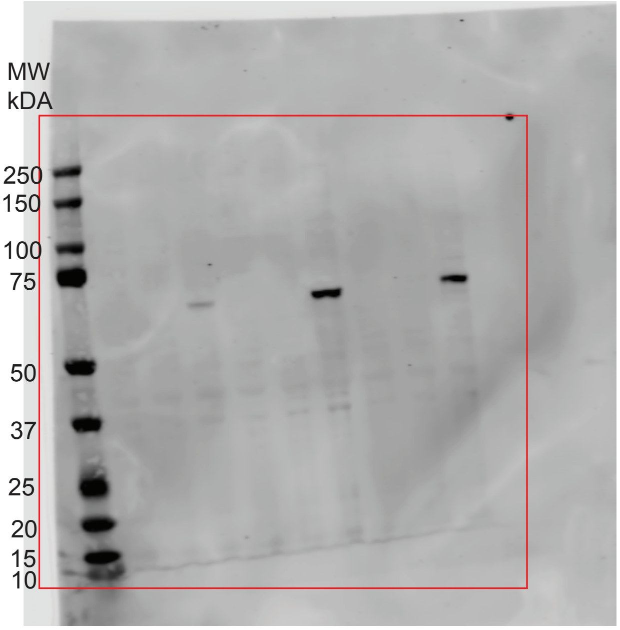


Anti-HA

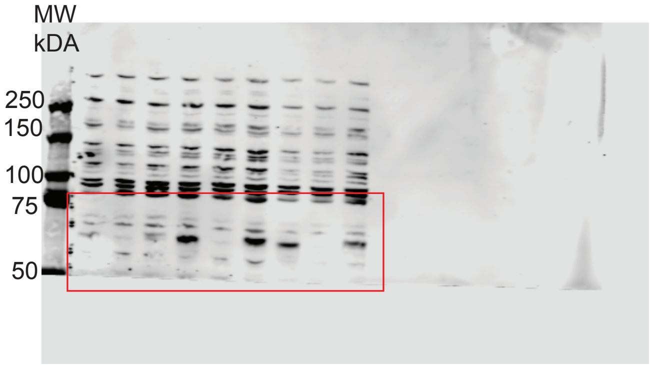


Anti-SP140

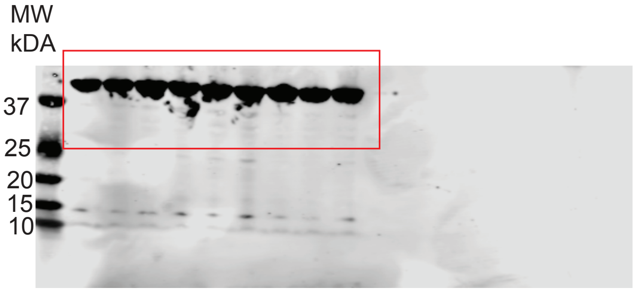


Anti-ACTIN (loading control for anti-SP140 blot, sample processing gel for anti-HA blot).
